# UNRAVELLING A HIDDEN SUBTYPE: MULTIOMICS REVEAL PSORIASIS-LIKE SIGNATURE WITH SURGICAL RELEVANCE IN A SUBSET OF PERIANAL FISTULIZING CROHN’S DISEASE

**DOI:** 10.64898/2026.07.25.740704

**Authors:** Saeed Abdurahiman, João Sabino, Sare Verstockt, Kristin Johnson, Lorenzo Giorio, Kaline Arnauts, Caitlin Van de Perre, Clara Caenepeel, Matthias Lenfant, Marc Ferrante, Tom Hillary, Anna-Teresa D’Hooghe, Gert De Hertogh, Manon E. Wildenberg, Christianne J. Buskens, Jeroen Raes, André D’Hoore, Séverine Vermeire, Gabriele Bislenghi, Bram Verstockt

**Author notes:** Corresponding author: Bram Verstockt, MD PhD Herestraat 49, 3000 Leuven BELGIUM.

## Abstract

**Background and aims:** Perianal fistulizing Crohn’s disease (pCD) affects 20% of patients with Crohn’s disease (CD) and severely impacts quality of life. Current therapies fail to provide sustained relief, and the molecular underpinnings of pCD remain poorly understood. This study aimed to elucidate the molecular landscape of perianal fistulas through multiomic profiling.

**Methods:** Paired fistula-tract and adjacent rectal mucosal biopsies were collected from 63 patients (48 with pCD, 15 with cryptoglandular fistula [CPTGL]). Longitudinal sampling generated 101 unique molecular profiles, comprising of both RNA sequencing (RNA-seq) and 16S rRNA sequencing from both tissue sites, followed by integrative multiomics and gene network analyses.

**Results:** Unsupervised clustering revealed three patient clusters primarily defined by host gene expression, with minimal contribution from microbial profiles. Fistulae in clusters 1 and 2 showed strong immune activation and epithelial–mesenchymal transition (EMT). In contrast, cluster 3 fistulae displayed keratinization and metabolic reprogramming resembling psoriatic skin, together with reduced JAK–STAT signalling. Cluster 3 was enriched for patients classified as TOpClass:2a, who are more suitable for surgical repair (p = 0.02). Conversely, cluster 2, characterized by rectal keratinization and EMT in both fistula and rectum, showed the highest MRI inflammatory-mass score (p = 0.02) and a greater risk of subsequent ileostomy (Kaplan–Meier; p = 0.008). An independent RNAseq dataset validated the keratinization signature in a subset of pCD fistulae.

**Conclusion:** Integrated multiomic analysis identified distinct molecular subtypes of pCD with surgical and therapeutic relevance. These findings refine the molecular understanding of pCD and support a precision medicine approach.

**What You Need to Know?:** *BACKGROUND AND CONTEXT:*

- Perianal fistulas affect 1 in 5 Crohn’s disease patients and have a severe negative impact on the quality of life of the patients.
- Most advanced IBD therapies are not efficacious for perianal Crohn’s disease.
- Surgical repair is not suitable for all patients.

*NEW FINDINGS:*

- This study is the first to delineate molecularly defined patient subtypes in perianal fistulizing Crohn’s disease. It further characterizes a psoriasis-like keratinization program in a subset of fistulas. Finally, it identifies molecular features and histological indicators that may explain-and potentially help predict-favorable surgical outcomes in selected patients

*LIMITATIONS:*

- Functional and mechanistic studies will be required to further dissect the drivers and dynamics of the epithelial remodeling identified.

*CLINICAL RESEARCH RELEVANCE:*

- Raises the possibility of refining of existing clinical stratification using molecular markers for improved therapeutic management and outcome.

*BASIC RESEARCH RELEVANCE:*

- Proposes keratinization as an opposing molecular process to EMT within the perianal fistula tract.
- Identifies robust gene signatures associated with EMT and keratinization in the fistula for further experimental and clinical studies.

## INTRODUCTION

Crohn’s disease (CD)-associated perianal fistula (pCD) affects about 1 in 5 patients with CD forming abnormal tracts extending from the anorectal mucosa to the perianal epidermis.^1–5^ Clinically, pCD often causes localized pain, discharge, faecal incontinence, and discomfort, limiting daily activities^2,6^ and significantly impacts the patients’ quality of life, influencing social, professional and emotional well-being.^7^ Even with a combined surgical and medical approach with infliximab as the gold standard,^8^ the treatment outcomes remain suboptimal, with about 50% of patients achieving fistula healing and only 36% sustained the response at one year.^9–11^ As a result, many pCD patients are at risk for defunctioning ostomy and/or proctectomy.^12^ Hence, there is an urgent need to better understand the underlying biological mechanism to enhance treatment efficiency and to identify novel therapeutic targets.

Proctitis and anorectal strictures have been strongly associated with fistula formation and persistence.^13^ Molecular profiling of rectal tissue corroborates these observations by demonstrating goblet cell dysfunction^14^ and a dysregulated myeloid-stromal dialogue, a phenomenon also seen in CD associated ileal fibrosis.^15,16^ Complementing these findings, Cao et al. reported parallel upregulation of interferon-gamma (IFN-γ) and Tumour necrosis factor-alpha (TNF-α) signalling in both fistula tracts and the ileum of pCD patients pointing to host intrinsic mechanisms underlying fistula pathogenesis.^17^

At the cellular level, epithelial–mesenchymal transition (EMT) has been proposed as a potential mechanism underlying fistula development, supported by the histological observation of transitional epithelial-mesenchymal cells lining the fistula tracts.^18^ Additional evidence comes from the presence of matrix metalloproteinases (MMP) expressing fibroblasts in fistula tract microenvironment. These fibroblasts produce matrix MMPs and collagens leading to an altered ECM structure.^19,20^ Gudino et al. further demonstrated TNF-like ligand 1A (TL1A)-Lymphotoxin-β axis as the mediator, independent of TNF, imprinting a potentially pro-fistula microenvironment in the rectum by modulation of epithelial cells and fibroblasts.^21^

Beyond host-intrinsic mechanisms, clinical improvement following treatment with antibiotics or faecal diversion also implicate a potential role for the microbiota in pCD.^22^ A microbial signature with reduced butyrogenic potential was found to be associated to perianal disease among patients with paediatric CD.^23^ Significant differences in the microbial composition exist also between matched stool and fistula tract.^24^ Overall, there are limited number of studies exploring the microbial composition in the fistula tract let alone the mechanistic links with the host tissue.^25^

Most perianal fistulas result from anal gland infections and are classified as cryptoglandular (CPTGL), distinguished from pCD.^26–28^ In contrast, pCD is thought to result from chronic transmural inflammation in CD, driven by a multifactorial interplay of genetic susceptibility, dysregulated immune responses, impaired tissue remodelling, and altered microbiota.^29,30^ While these factors have been implicated, a precise pathophysiology and a comprehensive molecular landscape of pCD, particularly the specific contributions and interactions between tissue microenvironment and microbiota, remains poorly characterized. To address this gap, we conducted a multiomic analysis of microbiota composition and host gene expression patterns in paired fistula and adjacent rectal tissues from both pCD and CPTGL patients. Our findings provide novel insights into the molecular mechanisms underpinning pCD, revealing not only potential novel therapeutic targets but also highlighting the patients most suitable for surgical repair.

## METHODS

### Ethics statements and approval

This study was conducted at the University Hospitals Leuven (Belgium). Consecutive patients with perianal disease, with or without CD, undergoing routine anal exploration under anaesthesia were prospectively recruited. All patients provided written informed consent for inclusion in the IBD Biobank of the University Hospitals Leuven, approved by the Institutional Review Board (S53684, S71238), and consented for publication.

### Sample Collection

Two paired biopsies were collected from fistula and rectum (adjacent to fistula orifice) of patients using an endoscopic forceps with CD-related fistula and CPTGL fistula during anal exploration by 2 highly specialised IBD colorectal surgeons. One pair was snap-frozen for microbial analysis, and another preserved in RNAlater for transcriptomics. For those patients who had magnetic resonance imaging (MRI) performed as part of routine clinical care within the 12 weeks prior to sampling and without any change in (medical) therapy, MRI findings were evaluated using the MAGNIFI–CD scoring system, specifically the inflammatory mass sub–score (range 0-5)^31^. Higher values indicate more active and severe disease.

### Bulk Transcriptomics

Biopsies in RNAlater were stored at −80°C until RNA extraction using the AllPrep DNA/RNA Mini kit (Qiagen, Germany). Libraries were generated following the TruSeq Stranded mRNA protocol (Illumina, USA) and sequenced on an Illumina HiSeq4000 platform. Reads were aligned to GRCh37 using Hisat2 (v2.1.0), and raw counts were obtained with HTSeq. Low-coverage samples (<1 million reads) were excluded, and only protein-coding genes (Ensembl GRCh37) with ≥5 counts in at least 20% of samples were retained for downstream analysis. Batch effect was assessed using BatchQC R package, and a batch correction was performed with Combat-Seq prior to MOFA/WGCNA.^32–34^

### Differential Gene expression Analysis

DESeq2 package was used for differential gene expression analysis, in which batch effects were modelled as a covariate.^35^ For Gene Set Enrichment Analysis (GSEA), we generated a ranked gene list based on log fold-change values and used clusterProfiler with Gene Ontology (GO) annotation.^36–38^

### Multiomic data integration

For multiomic factor analysis (MOFA), each patient-date instance with at least two of the four data types was treated as a unique profile. A total of 19 patients (17 CD, 2 CPTGL) contributed samples at more than 1 instance (Supp. Fig. 1A, Supp. Fig. 2A). No individual contributed more than one sampling per date, so clustering was performed on samplings (profiles) rather than individuals (See supplemental methods).

**Figure 1:**
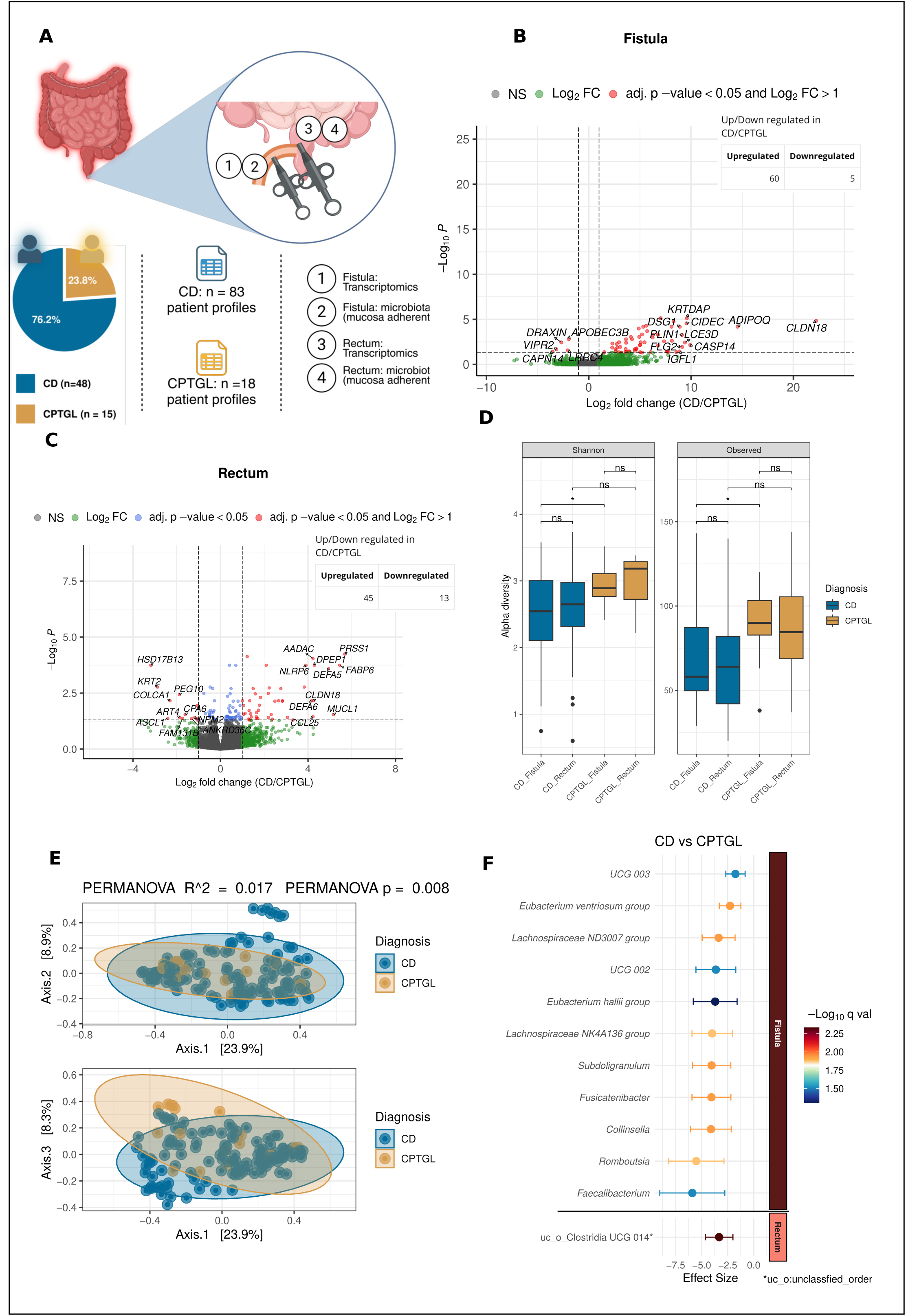
Transcriptomic and microbial alterations in fistula and rectal sites between Crohn’s disease and cryptoglandular fistula Comparison of transcriptomics profiles between CD and CPTGL. **A)** Scheme depicting sample collection for RNA sequencing and 16 rRNA sequencing assays. **B)** Volcano plot depicting DEGs in CD compared to CPTGL in the fistula. **C)** Volcano plot depicting DEGs in CD compared to CPTGL in the rectum. **D-E): Analysis of 16s rRNA sequencing data from CD and CPTGL D)** Alpha diversity metrics (Shannon index and Observed diversity) across the fistula and rectum in CD and CPTGL. **E)** Principal Coordinate Analysis plots depicting PERMANOVA test between CD and CPTGL in fistula. Axes 1-2 (top) and axes 2-3 (bottom). **F)** Differential taxa based on 16s microbial profiling in CD compared to CPTGL. Results are from MaAslin2 analysis. Fistula (top) takes into account Age and Sex. Rectum (bottom) takes into account Age, Sex and inflammation.

**Figure 2:**
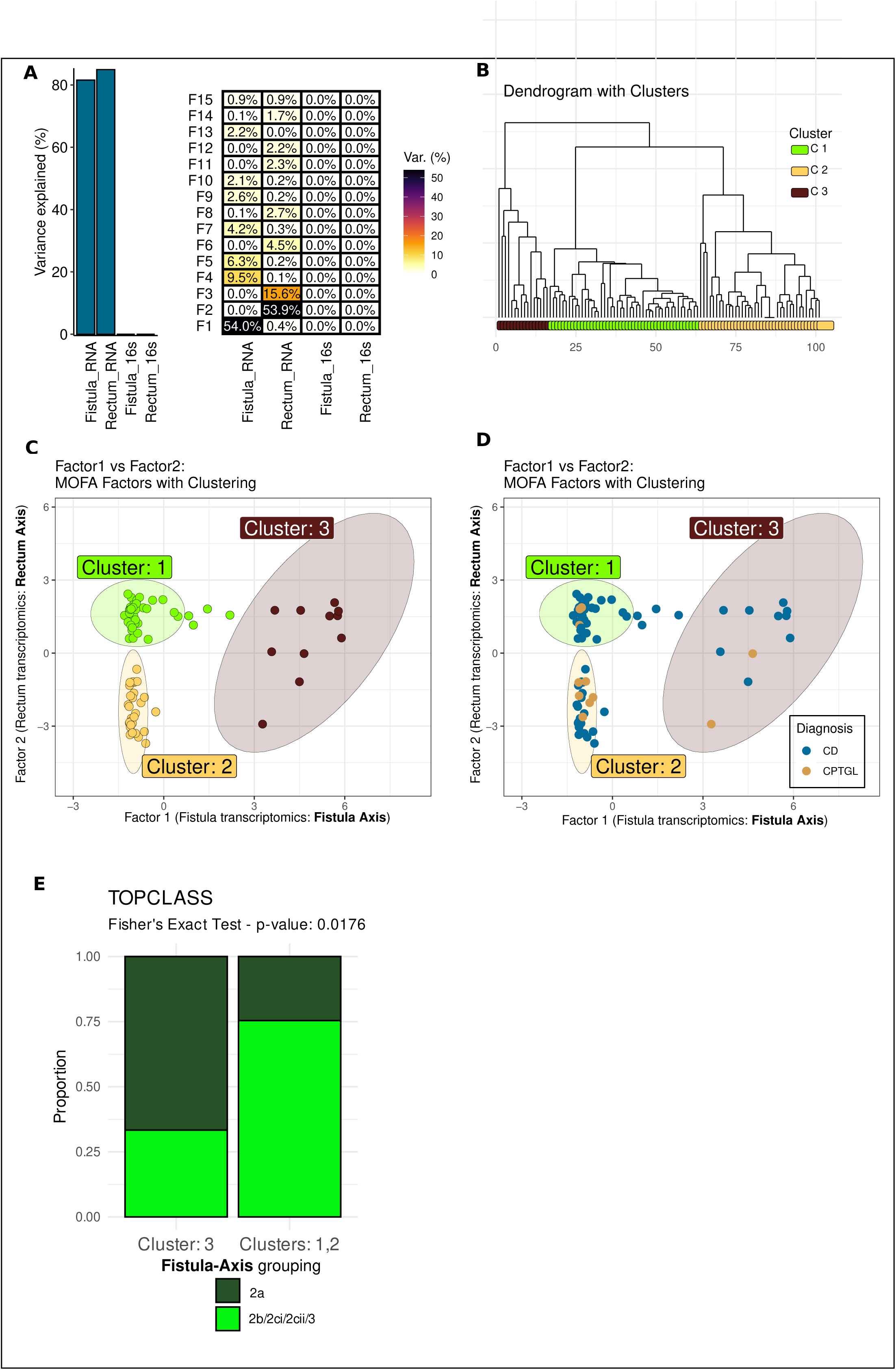
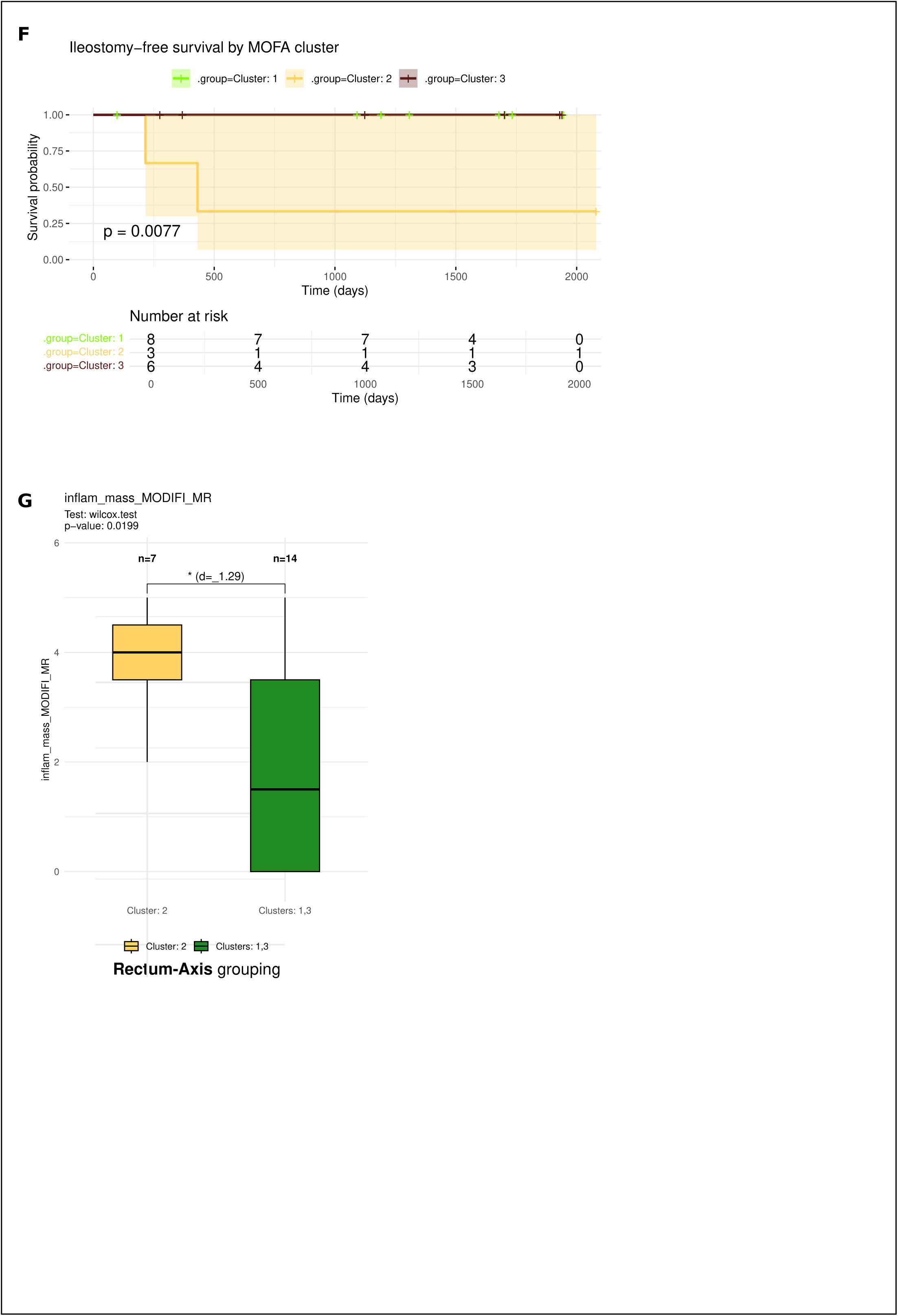
Three distinct molecular subtypes of fistulizing perianal disease Multiomics factor analysis (MOFA) and subsequent unsupervised clustering analysis. **A)** Barplot indicating percentage of variance explained by each omic layer in the MOFA model (left). Percentage of variance explained by each omic layer in each factor (right) **B)** Unsupervised hierarchical clustering of the patients based on the MOFA factors. **C-D**): Scatterplot depicting the clustering of patient profiles across MOFA Factor 1 (fistula axis) and MOFA Factor2 (rectum axis)**. C.** Scatterplot depicting the profiles across MOFA clustering. **D)** Scatterplot depicting the distribution of CD vs CPTGL profiles across the same rectum and fistula axes as in (C). **E)** TOpClass classification of patients across the two fistula groups (Fistula axis grouping). **F)** Kaplan-meier curve for the three clusters within patients classified as TOpClass 2a. **G)** Inflammatory mass sub-score from MAGNIFI-CD indexing across the two rectum axis groups.

For the input data, after batch correction using Combat-Seq, read counts were variance-stabilized via the VST function in DESeq2. Highly variable genes (HVGs) were identified in rectal and fistula transcriptomes using a custom iterative method: genes were ranked by variance, and an “elbow point” was defined geometrically to select HVGs. This process was repeated on non-HVGs to obtain a second HVG set. The procedure was applied separately to rectal, fistula, and combined datasets, and all HVG sets were pooled for MOFA input. For 16S data, a CLR transformation was applied. MOFA integrated all four datasets (Fistula_RNA, Rectum_RNA, Fistula_16S, Rectum_16S) using default settings with customizations: 15 factors, view scaling enabled, and 100 seeded initializations. The model with the highest ELBO was selected for downstream analysis.

### Weighted Gene Co-expression Network Analysis

Following batch correction and variance stabilization using VST from DESeq2, Weighted Gene Co-expression Network Analysis (WGCNA) was applied to identify gene co-expression modules.^28^ Optimal soft-thresholding power was selected using pickSoftThreshold to approximate scale-free topology. An adjacency matrix was computed and converted into a Topological Overlap Matrix (TOM), which was used for hierarchical clustering to generate a gene dendrogram. Modules were defined using dynamic tree cutting (minimum size: 30 genes) and merged if eigengene correlation exceeded 0.75. Hub genes were identified using ChooseTopHubGene, based on highest connectivity. Differences in module eigengene values between patient subgroups were assessed using Wilcoxon rank-sum or t-tests, depending on normality (evaluated via Shapiro-Wilk test)

### Single cell RNA-seq data re-analysis

Skin/psoriasis data was downloaded from GSE228421.^39^ Pre-processing of the data was done using Seurat (v5.1.0).^40,41^ Cells with less than 200 features were removed. Cells were annotated using CellTypist with predesigned model for adult skin as provided on CellTypist website based on Reynolds et al. (2021).^42–44^ The skin data was merged with Gut Data available from Kong et al. 2023.^45^ For Gut data, author annotation was available and hence used as such. When needed, the finer annotations were grouped into broader categories across organs. The top 10% percentile of most connected genes in the WGCNA module networks were extracted and used to create WGCNA module score for the single cell data using AddModuleScore function available in Seurat R package.^41^

### Functional analysis

GSEA or gene overrepresentation test were performed using ClusterProfiler (4.10.1) R package with gene annotations assembled from Molecular signature database.^34,46^

### Visualization

Visualizations were performed using ggplot2, ggpubr, cowplot, gridExtra R packages as well as using biorender.com.

## RESULTS

### Cohort description

In total 63 patients (48 CD, 15 CPTGL), contributed to a total of 101 (83 CD, 18 CPTGL) unique patient – date profiles, as some patients contributed at multiple timepoints during their disease journey (Fig 1A, Supp. Fig. 1A). Each patient-date profile was considered as a separate entity for multiomic analysis (see Methods) and henceforth referred to as ‘profile’ in this manuscript. Patient characteristics are summarized in Table S1 – S4. To minimize selection bias, all eligible patients were prospectively enrolled, regardless of treatment history. Prior drug and surgical exposures were recorded (Supplementary Table S1,S2). Demographic data were prospectively collected; specific features of perianal disease, including the TOpClass classification, were assessed by a multidisciplinary team (colorectal surgeons, gastroenterologists and radiologists), fully blinded to the molecular classification of the patients (Table S3).^39^ Of the 83 pCD profiles, 23 (27.7%) belonged to TOpClass 2a, while the rest were classified as 2b (56.6%), 2ci (8.4%), 2cii (3.6%) or 3 (3.6%). Proctitis was assessed by the surgeon during anal exploration and defined by a SES-CD > 2 in the rectal segment.

### Transcriptomic similarities between Crohn’s disease and cryptoglandular fistula

Biopsies were collected from the fistula tract and rectum (close to the internal fistula orifice) from patients with CD or CPTGL, as shown in Fig. 1A. Gene expression analysis of fistula tissue revealed only 60 up- and 5 downregulated (adj.p < 0.05, log2foldchange cut off = 1) genes in CD compared to CPTGL fistulas (Fig. 1B) with *CLDN18* being the most upregulated gene in CD (log2FC = 22.2, adj.p < 0.05). However, pathway analysis (GSEA) did not reveal any significantly enriched biological processes in any of the entities. A separate gene expression analysis in the rectal tissue, controlling for rectal inflammation, identified 45 up- and 13 downregulated genes (p < 0.05, log2foldchange cut off = 1) in CD compared to CPTGL (Fig. 1C). In summary, overall differences in the transcriptome were mainly driven by tissue origin (fistula vs rectum), rather than the underlying diagnosis (CD vs CPTGL) (Supp. Fig. 1B).

### Microbial alterations in fistula and rectal sites between Crohn’s disease and cryptoglandular fistula

In contrast, microbiome analysis showed significant differences in overall microbial diversity between CD and CPTGL fistulas (Shannon index, p < 0.05) (Fig. 1D). However, PCoA did not reveal distinct clustering between the two diagnostic groups (PERMANOVA R² = 0.017; Fig. 1E) or by TOpClass category (p > 0.05, R^2^ = 0.056) (Supp. Fig. 1C-E). Nevertheless, 11 bacterial genera were significantly reduced in CD fistulas compared to CPTGL (adj. p < 0.05), including *Faecalibacterium*, *Collinsella*, and *Romboutsia* (Fig. 1F), consistent with known dysbiosis in CD.^40–42^ In rectal biopsies, proctitis status was included as a covariate in the model; the only robust association observed was a reduced abundance of unclassified ASVs assigned to the Clostridia order in CD. Taken together, these data indicate that while overall diversity and community structure appear broadly similar, CD is characterized by loss of known beneficial taxa in the fistula compartment.

### Three distinct molecular subtypes of fistulizing perianal disease

To integrate transcriptomic and microbial findings in fistulizing perianal disease, we applied multiomic data integration across the four datasets: fistula and rectal RNA sequencing, as well as fistula and rectal microbiome data from both pCD and CPTGL.^34^ We considered each profile as independent as we did not observe systematic patient-patient similarity for the subset of patients with multiple sampling instances (see supplemental methods). This integrated multiomic analysis identified 15 “factors”, each representing a major source of biological variation across the datasets. These factors reflect coordinated biological signals that can help explain differences between patients, condensing complex multiomic data into clinically interpretable dimensions. Most of the observed variability was driven by transcriptomic data, while microbial composition contributed minimally (Fig. 2A). Unsupervised hierarchical clustering of integrated multiomic factors identified three distinct patient subtypes (Fig. 2B). These groups were driven mainly by two dominant transcriptomic components: one reflecting gene expression in fistula tissue (“fistula axis”) and the other reflecting gene expression in rectal tissue (“rectum axis”) (Fig. 2C). Together, these two axes explained more than half of the overall variation in the integrated dataset (Fig. 2A). When visualized in this two-dimensional framework, the fistula axis primarily separated Cluster 3 from Clusters 1-2, whereas the rectum axis distinguished Cluster 2 from Clusters 1 and 3 (Fig. 2C-2D). Because cluster assignment required information from both fistula and rectal gene-expression profiles, subsequent analyses were limited to profiles with transcriptomic data available from both tissue (70 pCD and 11 CPTGL). Given the clear tissue-specific structure captured by this integrative approach, these factors were used as the primary framework for downstream analyses.

Importantly, the molecular subtypes did not correlate with clinical diagnosis (CD vs. CPTGL; p=0.23), proctitis status (p=1.0), previous exposure to advanced biological therapy (p > 0.05), prior treatment with darvadstrocel (p=0.36), CD disease duration (p=0.36), time since first fistula (p=0.70) or history of a previous stoma (p=0.77) (Fig. 2D, Supp. Fig. 2B-M). No association was observed with perianal disease duration (p=0.38), arguing against these molecular patterns simply reflecting long-standing disease (Supp. Fig. 2N-O). In contrast, a clinically relevant association emerged with the TOpClass classification. Profiles from patients with a chronic symptomatic fistula considered suitable for fistula repair (TOpClass 2a) were overrepresented in Cluster 3 (p=0.02) (Fig. 2E). Because TOpClass 2a alone does not ensure an ileostomy-free course, we further examined prognosis within this subgroup. Notably, TOpClass 2a patients in cluster 2 had significantly poorer ileostomy-free survival compared with those in the other clusters (Fig. 2F).

For a subset of CD profiles (n=21, Table S5), pelvic MRI within a 12-week period prior to sampling was available. The MAGNIFI-CD inflammatory mass score (range 0 to 5) was significantly higher in Cluster 2 (p=0.02, Fig. 2G) and was numerically lowest in Cluster 3 (p=0.07) (Supp. Fig. 2P-Q). Consistently, inflammatory mass scores were significantly lower in TOpClass 2a compared to 2b (p=0.003) (Supp. Fig. 2R).^31^

Finally, given the minimal contribution of microbial data to the overall variation, differential abundance analysis^43^ did not identify any bacterial genera associated with the molecular subtypes (data not shown).

Overall, these findings suggest in pCD, host-gene expression patterns play a more prominent role than microbial composition in defining molecular subtypes and are linked to clinically meaningful outcomes.

### Two divergent molecular pathways in fistula formation

Because transcriptomic data largely drove clustering, we next sought to define the molecular features underlying the three patient subtypes. Weighted Gene Co-expression Network Analysis (WGCNA) was applied to identify groups of co-expressed genes (gene-modules), which often represent shared biological function.^32^

Fourteen gene modules were identified in the fistula tissue. Five showed strong correlations with the molecular clusters (|r| > 0.5, adjusted p < 0.05) (Fig. 3A, Supp. Fig 3A-B). Specifically, two gene-modules (Fistula_M6 and Fistula_M7) were significantly associated with Cluster 3 patient profiles (Fig. 3B, Supp. Fig. 3A-B). Fistula_M6 was enriched for oxidative phosphorylation and aerobic respiration, indicating increased metabolic activity (Fig. 3B; Supp. Fig. 3C). Module Fistula_M7 was enriched for keratinisation-related pathways, including skin development, keratinocyte differentiation and cornified-envelope formation (Fig. 3B; Supp. Fig. 3D). Together, these findings suggest that Cluster 3 fistulae exhibit an epithelial-like differentiation program, possibly reflecting an abnormal differentiation.

**Figure 3:**
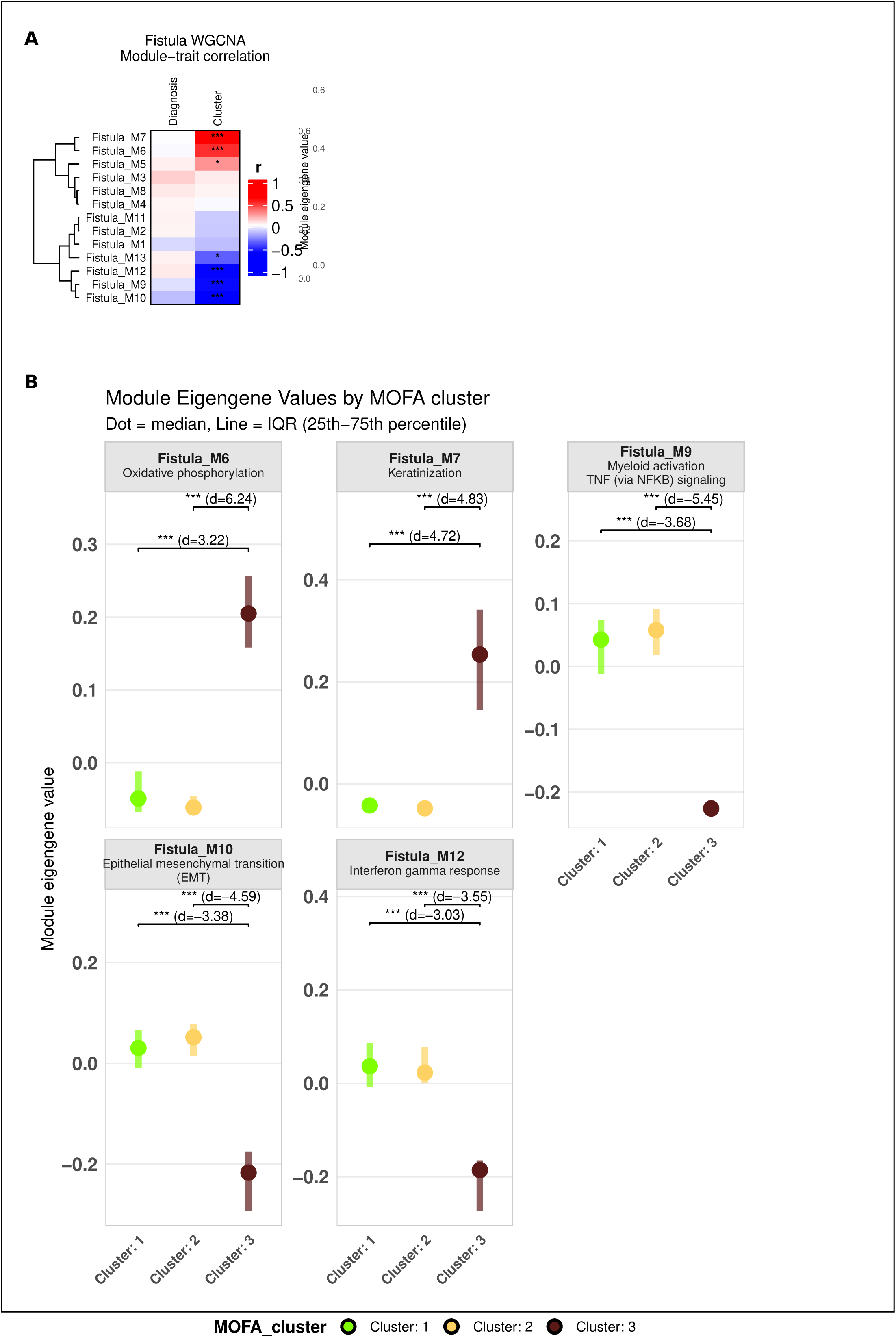
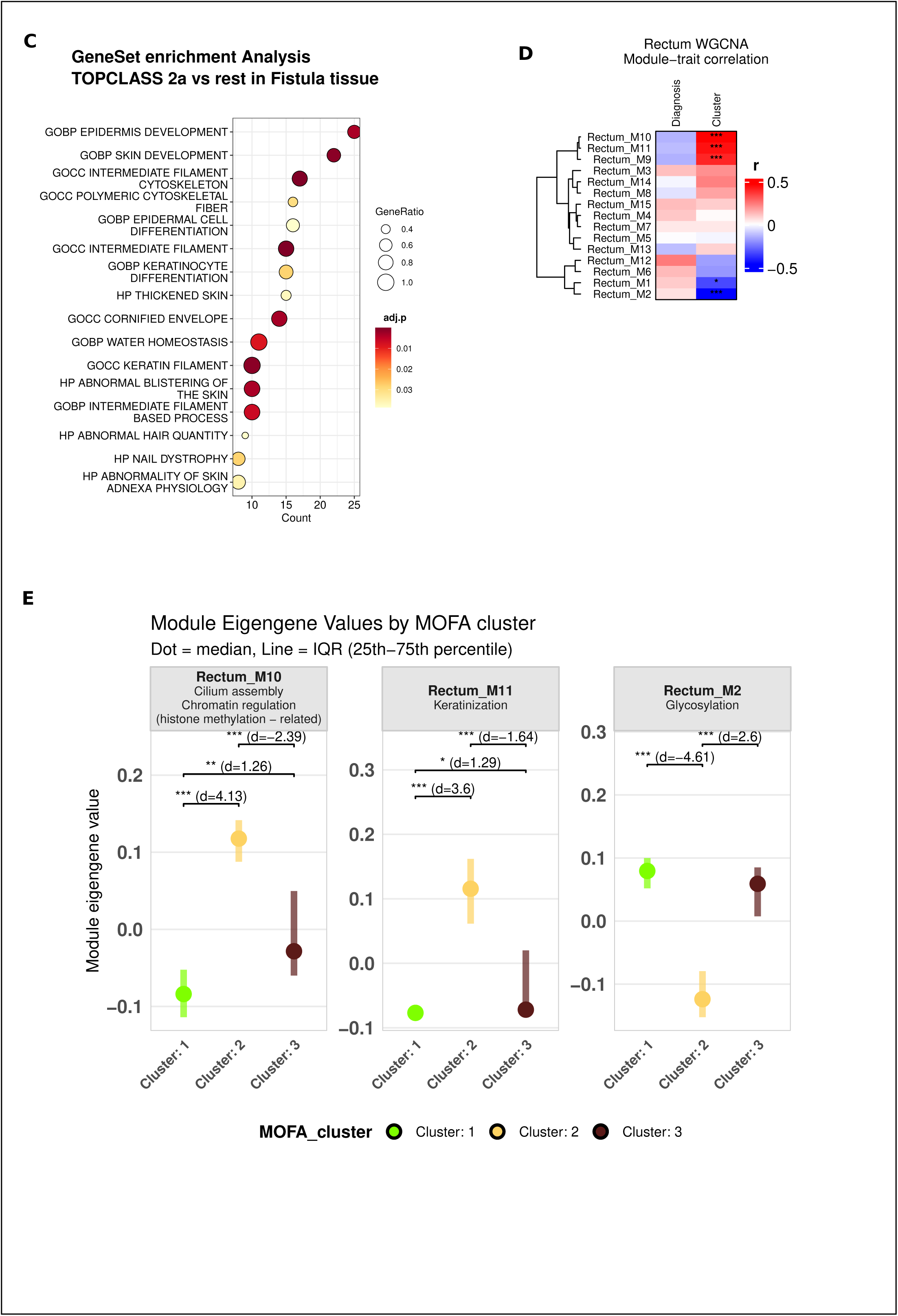
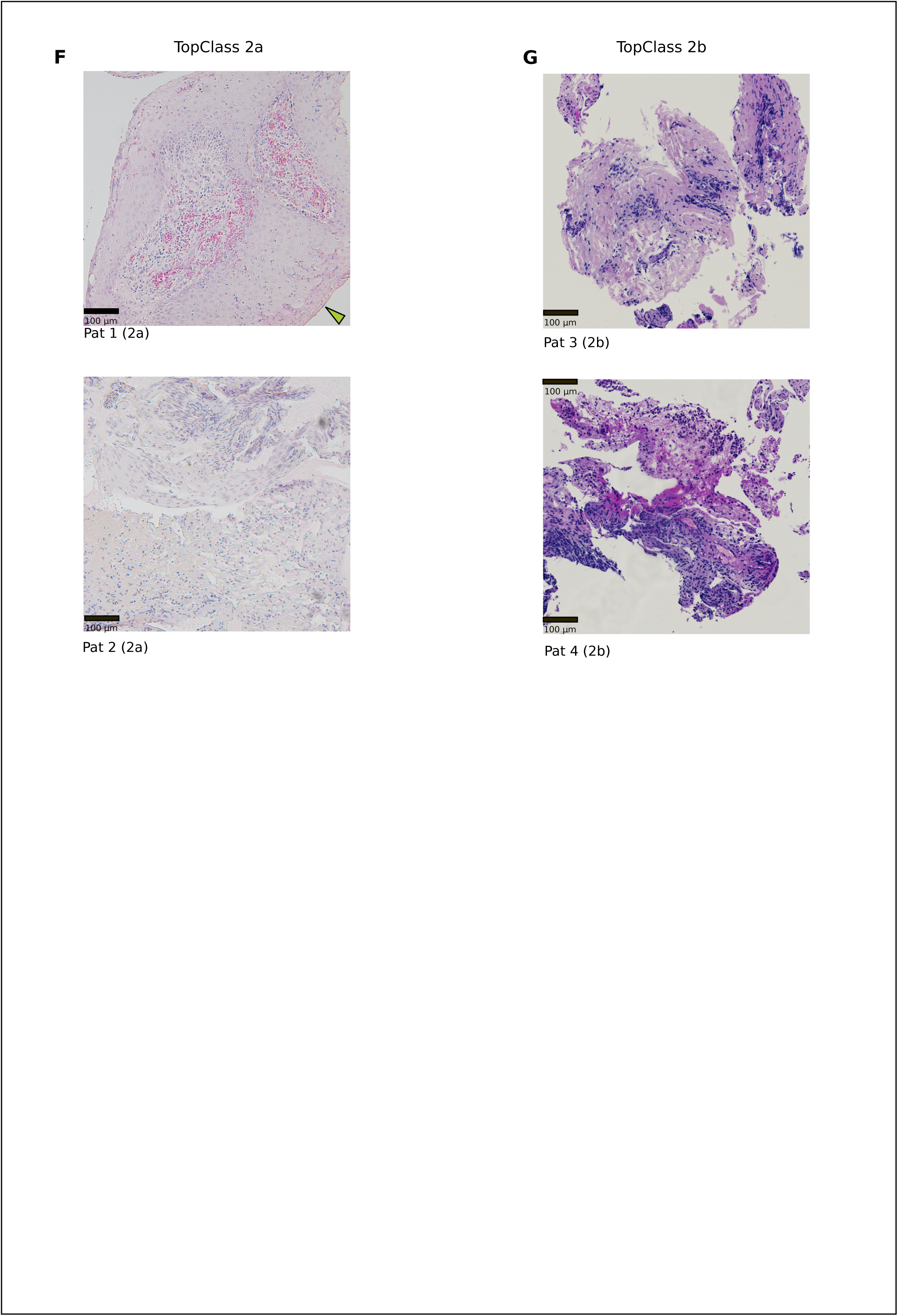
Divergent molecular pathways in fistula and rectum. **A)** Heatmap depicting correlation of WGCNA module eigengenes with the patient traits in fistula tissue. **B)** Module eigengene values for the five fistula gene modules with most significant correlations with the traits in **(A)** plotted across the MOFA clusters. For each module, key significantly enriched functional terms identified through enrichment analysis are highlighted within the corresponding panel. Effect size reported as standardized mean difference (Cohen’s d) indicated as ‘d’ **C)** Gene set enrichment analysis for the comparison between TOpClass 2a vs others. **D)** Heatmap depicting correlation of WGCNA module eigengenes with the patient traits in rectal tissue **E)** Module eigengene values for the three rectal gene modules with most significant correlations with the traits in (C) plotted across the MOFA clusters. For each module, key significantly enriched functional terms identified through enrichment analysis are highlighted within the corresponding panel. ffect size reported as standardized mean difference (Cohen’s d) indicated as ‘d’ **F)** HE images of fistula tract biopsies from **two** TopClass 2A patients (top and bottom from 2 patients). Green arrow indicate region depicting parakeratosis. **G)** HE images of fistula tract biopsies from two TopClass 2b patients (top and bottom from 2 patients).

In contrast, three gene-modules (Fistula_M9, Fistula_M10, Fistula_M12) were predominant in Clusters 1 and 2, and relatively reduced in Cluster 3 (Fig 3A-B, Supp. Fig. 3A-B). These modules were enriched for inflammatory and tissue-remodelling processes. Module Fistula_M9 reflected myeloid inflammatory responses and IFN-γ and TNF signalling (Fig 3B, Supp. Fig. 3E). Module Fistula_M10 was linked to collagen fibril organization and EMT, key pathways in tissue remodelling and fibrosis (Fig. 3B, Supp. Fig. 3F). Module Fistula_M12 was associated with lymphoid immune responses, particularly IFN-α and IFN-γ signalling (Fig. 3K-L, Supp. Fig. 3G). Differential gene expression analysis confirmed EMT activity in Cluster 3. Key EMT related transcription factors (*SNAI1*, *SNAI2* and *TWIST1*) and EMT-inducing cytokines (*TGFB1*, *IL1B*, *TNF*, *IL6* and *OSM*) were downregulated (adj.p < 0.05), while the EMT suppressor *GRHL2* was upregulated (adj.p < 0.05, Supp. Fig. 3H).^44,45^ Inflammatory signatures accompanied EMT, with markers of inflammatory monocytes and neutrophils upregulated in fistulas of Clusters 1 and 2. Notably, *S100A8* and *S100A9*, components of calprotectin, were also upregulated (adj.p < 0.05) in fistula of Cluster 3, coinciding with keratinization (Supp. Fig. 3H).

These findings indicate that fistulae in Clusters 1 and 2 are primarily driven by chronic inflammation and tissue remodelling, consistent with the classical understanding pCD. Cluster 3, which was enriched for TOpClass 2a, showed evidence of an epidermal-like differentiation program. Gene set enrichment analysis (GSEA) using genes differentially expressed in TOpClass 2a fistulae confirmed significant enrichment of this program (Fig. 3C).

Overall, the data suggest two distinct molecular pathways in perianal fistulizing disease: one characterised by inflammation and remodelling, and another marked by epidermal-like differentiation and enhanced metabolic activity, which might be linked to a greater likelihood of successful surgical repair.

In line with the molecular indications of epidermis development and keratinization, we observed squamous epithelium in fistulous tracts of TopClass 2a patients. Fig 4F illustrates squamous epithelium with occasional parakeratosis (green arrow) in the fistulous tracts of two TopClass 2a patients. Fig 4G illustrates highly inflamed granulation tissue or fibrotic tissue in fistulous tracts of two TopClass 2b patients.

**Figure 4:**
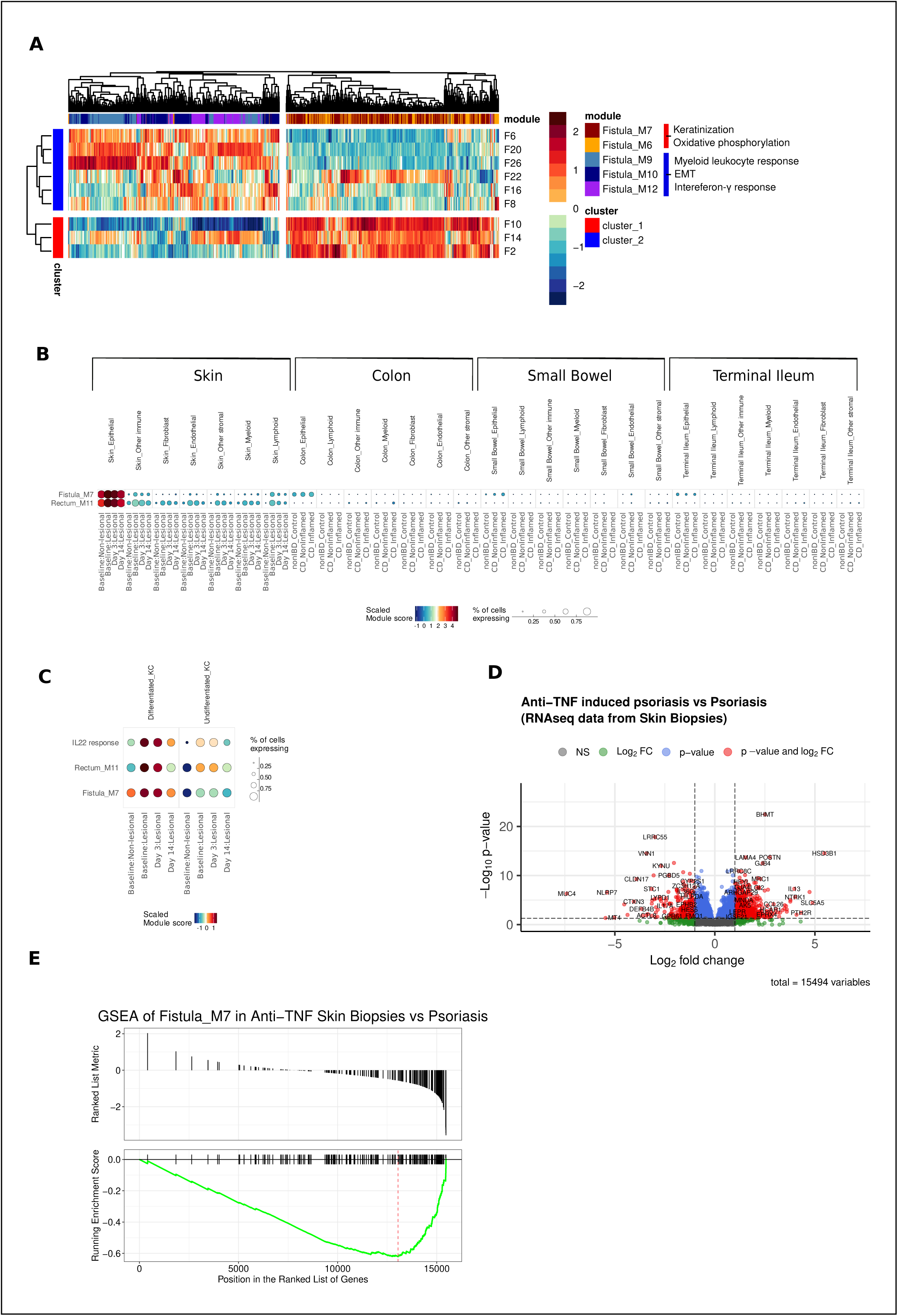
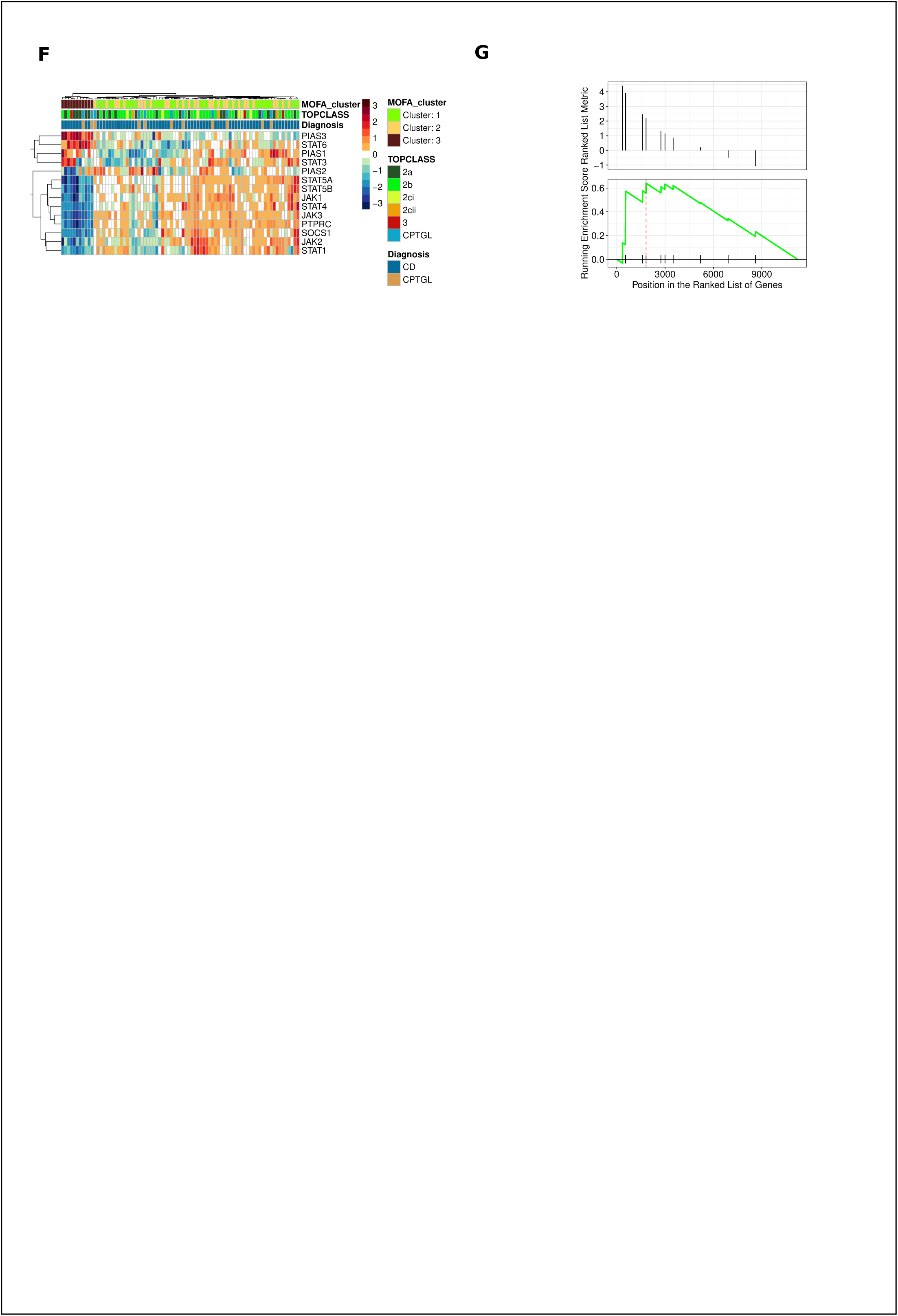
Validation of fistula subtypes and similarity to epidermal tissue. **A)** Heatmap depicting the genes of the fistula modules on external fistula gene expression data. Horizontal rows indicate patients (colored by patient cluster) and columns indicate genes (colored by fistula modules). **B)** Top hub genes (90^th^ percentile) from each of the keratinization related modules (Fistula M7, Rectum_M11) were used to create a gene module score in the combined single cell RNA sequencing data from psoriasis study (Francis et. al.) and CD (Kong et, al.) **C)** Subset of the psoriasis single cell data set comprising only keratinocytes. Module score computed as in (**A**) for keratinization modules and IL22 response genes (MsigDB: Human Gene Set: ZHENG_IL22_SIGNALING_UP) depicted specifically in the longitudinally profiled keratinocytes from psoriasis study at baseline and post IL23 treatment. **E**) Volcano plot depicting differential gene expression analysis of RNA seq with skin biopsies from anti-TNF induced psoriasis in IBD patients (n=7) vs idiopathic psoriasis (n=10). **G**) GSEA plot showing enrichment of Fistula_M7 keratinization module in psoriasis over anti TNF induced psoriasis in IBD patients (p<0.001) **F)** Heatmap showing genes in the JAK-STAT signaling gene set (Panther pathways) in the fistula tissue. **G)** GSEA analysis for JAK-STAT pathway in fistula (p value = 0.02).

### Distinct rectal gene expression patterns in patients with perianal disease

A similar analysis of rectal transcriptomic data revealed distinct molecular patterns that parallel the previously defined patient subtypes. Three rectal gene co-expression modules were associated with these clusters (Fig. 3D). Cluster 2 exhibited high expression of modules Rectum_M10 and Rectum_M11 (Fig. 3E, Supp. Fig. 3I-J). These modules were enriched for pathways related to epigenetic and post-transcriptional regulation (e.g., histone modification, RNA methylation) and keratinization (e.g., skin development, keratinocyte differentiation), respectively (Fig. 3E, Supp. Fig. 3 K-L, Supp. Material2). This indicates that rectal tissue in Cluster 2 also displays an epidermal-like gene expression signature, similar to that observed in fistula tissue in Cluster 3.

In contrast, module Rectum_M2, which was relatively reduced in Cluster 2, was linked to glycosylation and organic acid metabolism, suggesting metabolic differences between subtypes (Fig. 3D, Supp. Fig. 3M, Supp. Material2). Unlike the fistula-derived transcriptomic component, which clearly distinguished TOpClass 2a-enriched Cluster 3, the rectal transcriptomic patterns were not specifically associated with TOpClass categories (Supp. Fig. 3N). Proctitis status also did not correspond to the rectal molecular groupings (p=1.0) (Supp. Fig. 3O).

To further explore tissue-remodelling pathways, we examined expression of EMT-related genes in rectal tissue. EMT-transcription factors *SNAI1*, *SNAI2* and *TWIST1* were significantly upregulated in Cluster 2 rectal samples compared to Cluster 1 and 3 (adj.p < 0.05, Supp. Fig. 3P).

Overall, these findings suggest concurrent keratinization and EMT activity in the rectum of Cluster 2 profiles and highlight previously unrecognized molecular heterogeneity in rectal tissue among patients with perianal disease.

### Analysis of external data validates the existence of the distinct subtypes

To externally validate these findings, we analysed a previously published RNA-sequencing dataset from fistula tract samples.^46^ Unsupervised hierarchical clustering clearly separated patients into two distinct groups (approximately unbiased (p-value ∼ 100% by multiscale bootstrap resampling) (Supp. Fig. 4A).^47^ Next, we repeated the clustering using only genes representing the two divergent molecular pathways programs identified in our cohort: immune activation and EMT (Fistula_M9, Fistula_M10, Fistula_M12), and keratinization coupled with oxidative phosphorylation (Fistula_M7, Fistula_M6). This gene-restricted analysis produced patient groupings identical to the initial unsupervised groupings (adjusted Rand index = 1; permutation test, p < 0.05; data not shown) (Fig. 4A, Supp. Fig. 4A). Together, these results confirm the presence of two fundamentally distinct molecular profiles within perianal fistula tissues.

### A psoriasis-like molecular signature in perianal fistula

To further investigate the ectopic, epidermal-like gene expression signature observed in one molecular subtype, we examined this signature in publicly available single-cell RNA sequencing (scRNA-seq) datasets from psoriasis and IBD patients.^48,49^ Publicly available scRNA-seq datasets from perianal fistulas are limited to very small cohorts (n = 3 patients) and either capture inadequately fistula tissue or lack epithelial cells, making them unsuitable for evaluating keratinization signatures (data not shown).^16,17^

However, the keratinization-associated gene modules identified in our study (Fistula_M7 and Rectum_M11) were strongly enriched in skin epithelial cells, rather than gut-specific cell populations. In particular, they were prominent in keratinocytes from psoriatic lesions compared to non-lesional skin (Fig. 4B, Supp. Fig. 4B).^48^ These signatures were reduced in lesional keratinocytes following anti-IL23 treatment, with a similar pattern observed for the IL-22 pathway (Fig. 4C). The gene *GLTP* emerged as a central hub within the Fistula_M7 keratinization module (Supplementary Material 3) and has been reported among the most differentially expressed proteins in psoriatic lesions compared with non-lesional skin.^50,51^

Together, these findings indicate that a subset of perianal fistulas exhibits a molecular program that more closely resembles psoriatic keratinocytes than intestinal inflammatory tissue, suggesting an ectopic epidermal differentiation process, including features such as parakeratosis characteristic of skin disorders such as psoriasis.^52^

### Potential clinical and therapeutic implications

Patients with a Cluster 3 molecular profile may be particularly suitable for surgical repair, consistent with the enrichment of TOpClass 2a in this group. The psoriasis-like gene signature observed in Cluster 3 fistula tissue was also shown to be modulated by IL-23-targeted therapy in psoriatic skin (Fig. 4B, Supp. Fig. 4B), suggesting that anti-IL23 therapy could represent a therapeutic option for these patients. To determine whether the psoriasis-like signature we observed reflects anti-TNF-induced psoriasis - reported in a subset of patients with inflammatory bowel disease (IBD) - we compared transcriptomic profiles from conventional psoriasis skin lesions (non-IBD) with those from anti–TNF-induced psoriasis– like lesions in IBD patients.^53^ We found that the top genes within the fistula keratinization module were significantly enriched in conventional psoriasis lesions compared with anti– TNF-induced lesions. This suggests that the keratinization signature identified in fistula tissue may not be an ectopic representation of anti–TNF-associated pathology. (Fig 4D-E)

To explore whether other targeted treatment pathways aligned with the molecular subtypes, we assessed genes involved in the JAK-STAT signalling. Majority of JAK-STAT pathway genes were significantly upregulated (adj.p < 0.05) in the fistula tissue from Clusters 1 and 2 compared to Cluster 3, and hierarchical clustering based on these genes clearly distinguished Cluster 3 (Fig. 4F).^54^ Gene set enrichment analysis confirmed significantly higher JAK-STAT pathway activity in Clusters 1 and 2 relative to Cluster 3 (p=0.02) (Fig. 4G). Although *STAT3*, *STAT6*, and IL-4/IL-13 receptor genes were elevated, core JAK kinases (JAK1, JAK2, JAK3, TYK2) were significantly reduced in Cluster 3 fistulae (Supp. Fig. 4C-D). No clear enrichment for this pathway was observed in rectal tissue (data not shown). ^60^ Overall, these results indicate that different molecular subtypes of pCD may respond differently to targeted therapies, supporting a more individualised treatment approach.

## DISCUSSION

Although the TOpClass clinical classification offers a crucial framework for managing pCD, it remains insufficient for personalized therapy due to limited insights into the underlying biology.^39,55^ In this study, we assembled the largest multiomic dataset of perianal fistula to date, comprising of 101 molecular profiles with paired microbial and transcriptomic data from fistula tracts and adjacent rectal tissues, enabling a comprehensive molecular characterization.

Supervised transcriptomic analysis mirrored earlier reports indicating minimal differences between CD-associated and CPTGL fistulas, implying a shared molecular phenotype despite potential divergent etiologies.^27,28,56,57^ The modest molecular differences observed despite their contrasting clinical courses are most likely attributable to persistent inflammation in CD, in contrast to CPTGL, as demonstrated by Becker et. al.^57^ In contrast, microbial profiles showed variation based on molecular assay, particularly in CD fistulas, where reductions in beneficial taxa resembled the dysbiosis commonly observed in luminal CD.^58^

Unsupervised integrative analysis revealed three molecular subtypes that did not align with conventional clinical variables such as diagnosis, disease duration, prior biologic exposure or proctitis status. This previously unrecognized biological heterogeneity may help explain why current treatment strategies, which rely mainly on clinical features, often yield suboptimal outcomes.

Although healing of proctitis is a central therapeutic goal, particularly prior to surgical repair, the rectal molecular programs identified here did not segregate by intra-operative proctitis status. Instead, variation was driven primarily by epithelial differentiation and glycosylation-associated pathways, with an EMT-like signature in one cluster. These data suggest that clinically apparent proctitis may not fully capture repair-relevant mucosal states, and that non-inflammatory remodeling/barrier programs may persist despite absence or resolution of proctitis

Functionally, Clusters 1 and 2 were characterized by strong inflammatory, immune and EMT signatures, consistent with the traditional view of fistulizing CD.^25,59^ In contrast, Cluster 3 profiles, enriched for clinically repairable TOpClass 2a fistulas, displayed prominent keratinization, epidermal differentiation, and metabolic gene signatures, suggesting a distinct pathogenic pathway resembling epidermal disorders. These divergent programs were validated in an external RNAseq dataset.^46^ Notably, Cluster 2, characterized by rectal keratinization with EMT in both fistula and rectum, showed the most severe imaging features. Patients in this cluster classified as TOpClass 2a nonetheless had a significantly higher risk of ileostomy than those in Clusters 1 and 3, indicating that transcriptomic markers, particularly simultaneous keratinization and EMT in the rectum, could refine current clinical classification and guide management.

Earlier histological studies reported epithelialization of fistula tracts in a subset of patients,^60^ while more recent single-cell and spatial transcriptomic analyses identified these cells as squamous epithelial cells of unknown origin.^17,57^ Indeed, we also observed squamous epithelium in fistulous tracts of TopClass 2a patients that we saw enriched in Cluster 3.

The presence of squamous epithelial cells accounts for the keratinization and high expression of fecal calprotectin genes (S100A8/A9) observed in the Cluster 3 subset. Becker et al. demonstrated that, in fistula tissue, S100A8/A9 can localize to the nucleus, where it may function as a transcriptional regulator rather than being secreted as calprotectin, as typically seen in luminal CD.^57^ Therefore, the apparent upregulation of calprotectin transcripts (S100A8/A9) in Cluster 3 fistula tissue should not be interpreted as evidence of neutrophil-driven inflammation. This aligns with our observation that Cluster 3 fistulae display a keratinisation-dominant, low-EMT, low-Inflammation phenotype despite high S100A8/A9 transcript expression.^17,57,60^

Therapeutically, our proposed molecular stratification reveals distinct treatment opportunities. The psoriasis-like epidermal differentiation program observed in Cluster 3 fistula and Cluster 2 rectum resembles pathways responsive to IL–23 blockade, supporting exploration of anti-IL-23 therapies in these patients.^61–63^ Anti-IL23 therapy has recently been tested in pCD in the FUZION trial with guselkumab (NCT05347095). Despite any molecular pre-screening, the primary endpoint at week 24 was met, demonstrating the clinical relevance of anti-IL23 therapy in pCD for fistula closure.^64^ Conversely, Clusters 1 and 2 showed an upregulation of JAK-STAT pathway activity, providing plausible biological context for the observed efficacy of upadacitinib, a JAK1 inhibitor, in some pCD patients.^65^ Together, these data support a precision-medicine approach in which treatment selection reflects molecular subtypes, as well as clinical stage.

This study has some limitations. Its cross-sectional and exploratory design precludes causal inference, although sampling was standardised and performed by two highly IBD-specialised colorectal surgeons. Prior surgeries and immunomodulatory treatments may influence tissue gene expression, but clustering was not associated with prior drug exposure or disease duration. At present, how these molecular states evolve in response to specific surgical approaches and medical therapies remains unknown. Longitudinal sampling before and after protocolized interventions will therefore be essential to determine whether treatments shift tissue programs toward more favorable states and whether such shifts mediate improved healing and long-term outcomes. Regarding host–microbe interactions, 16S rRNA profiling captured known differences between CD and non-CD patients, but did not parallel the strong host transcriptomic subtypes. This likely reflects limited microbial biomass and the taxonomic resolution of 16S sequencing in tissue biopsies rather than absence of relevant interactions. Deeper metagenomic approaches may better define microbial contributions.

Leveraging the largest transcriptomic cohort reported to date in pCD, this study integrates and extends recent molecular findings to show that keratinization and EMT represent distinct and divergent cellular programs within fistulizing disease. Crucially, these molecular programs were linked to clinically meaningful differences in disease severity and prognosis, demonstrating that molecular state captures biologically relevant information beyond conventional clinical stratification. By defining putatively favorable and unfavorable molecular states across rectum and fistula tissue, our work provides a conceptual and analytical framework for future mechanistic and translational studies, including biomarker discovery and molecularly guided therapeutic strategies. Ultimately, identifying actionable determinants of molecular state may open avenues to modulate fistula biology directly, with the potential to enhance the effectiveness of both surgical and medical interventions in pCD through more precisely targeted treatment approaches.

In conclusion, these findings reveal substantial and previously unrecognized molecular heterogeneity in perianal CD, and identify an ectopic epidermal-like differentiation program resembling psoriatic skin. This challenges the traditional view of CD fistulas as uniformly inflammatory and highlights distinct pathogenic pathways with potential therapeutic implications. Further functional validation will be required, but these data provide a foundation for precision-medicine strategies aimed at improving outcomes in complex perianal disease.

## ACKNOWLEDGEMENTS

We express our gratitude to all lab technicians (Helene Blevi, Tamara Coopmans, Hannelore Hoogsteyn, Sophie Organe, Margot De Smet and Maja Tubak), our IBD biobank managers (Vera Ballet and Justien Degry) and IBD Leuven lab manager, Dr. Arno Cuvry, for their work in coordinating, collecting and processing the patient samples. We also thank the Genomics Core at UZ/KU Leuven for their technical support. We thank Dr. Padhmanand Sudhakar for support.

Sare Verstockt was a postdoctoral fellow of the Research Foundation – Flanders (FWO, Belgium). Kristin Johnson is supported by Swedish governmental funding of clinical research (ALF) and received grants from Maggie Stephens foundation, Gastroenterological Research Fund Sweden, the Royal Physiographic Society of Lund, Medical Society in Lund, Olle Olsson foundation and Swedish Society of Radiology. João Sabino and Marc Ferrante serve as Senior Clinical Researchers for the FWO, while Séverine Vermeire held a BOF-FKO grant from KU Leuven. Bram Verstockt is supported by the Clinical Research Fund (KOOR) of the University Hospitals Leuven and by the KU Leuven Research Council. The resources provided by the VSC (Flemish Supercomputer Center), funded by the Research foundation – Flanders (FWO) and the Flemish Government was utilized in this work.

## AUTHOR CONTRIBUTIONS

SA, SéV, GB, BV conceptualized the study. SA, JS, KJ, GB, BV devised methodology. SA provided software. SA performed formal analysis with support from JS. SA, JS, SA, JS, SaV, KJ, LG, KA, CVP, CC, ML, MF, MW, CB, AD, GB, GDH and BV performed investigation. JR, SéV, BV provided resources. JS, ML, GB, SaV, BV curated data. SA performed visualization. SéV and BV acquired funding. BV managed the project. SA and BV wrote the original draft. All authors reviewed and edited the manuscript.

## DATA AVAILABILITY

Data will be made available upon request from the corresponding author.

## PATIENT AND PUBLIC INVOLVEMENT

Neither patients nor the public were involved in the design, execution, reporting, or dissemination of this research.

---

**Supplementary figure 1:**
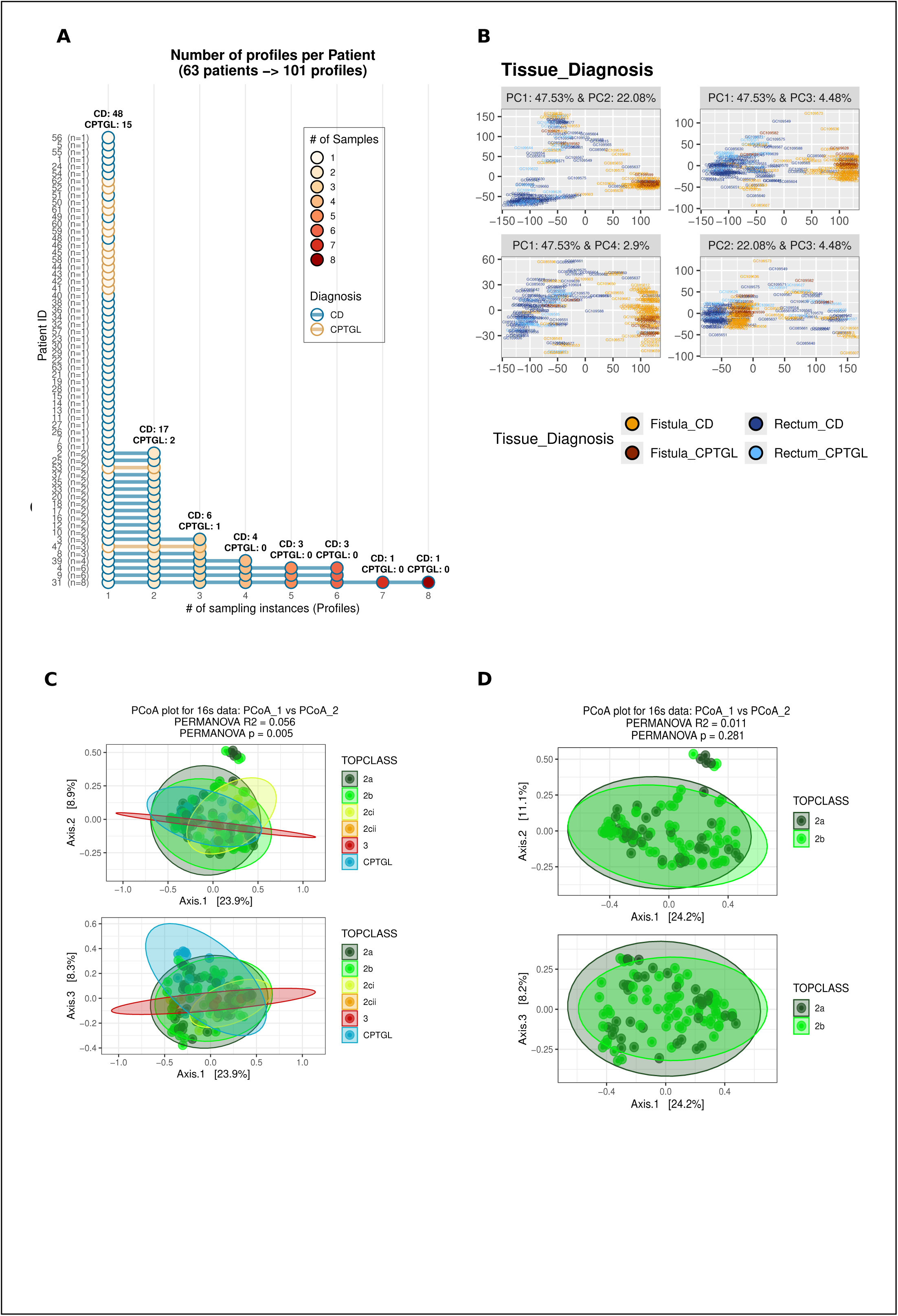
**A)** Plot showing the number of profiles each patient contributed (each row is 1 patient). **B)** Principal component analysis of the entire RNA seq data set of 178 samples from fistula and rectum of CD and CPTGL patients. **C-D)** PCoA plot and PERMANOVA analysis depicted according to TOpClass classification.

**Supplementary figure 2:**
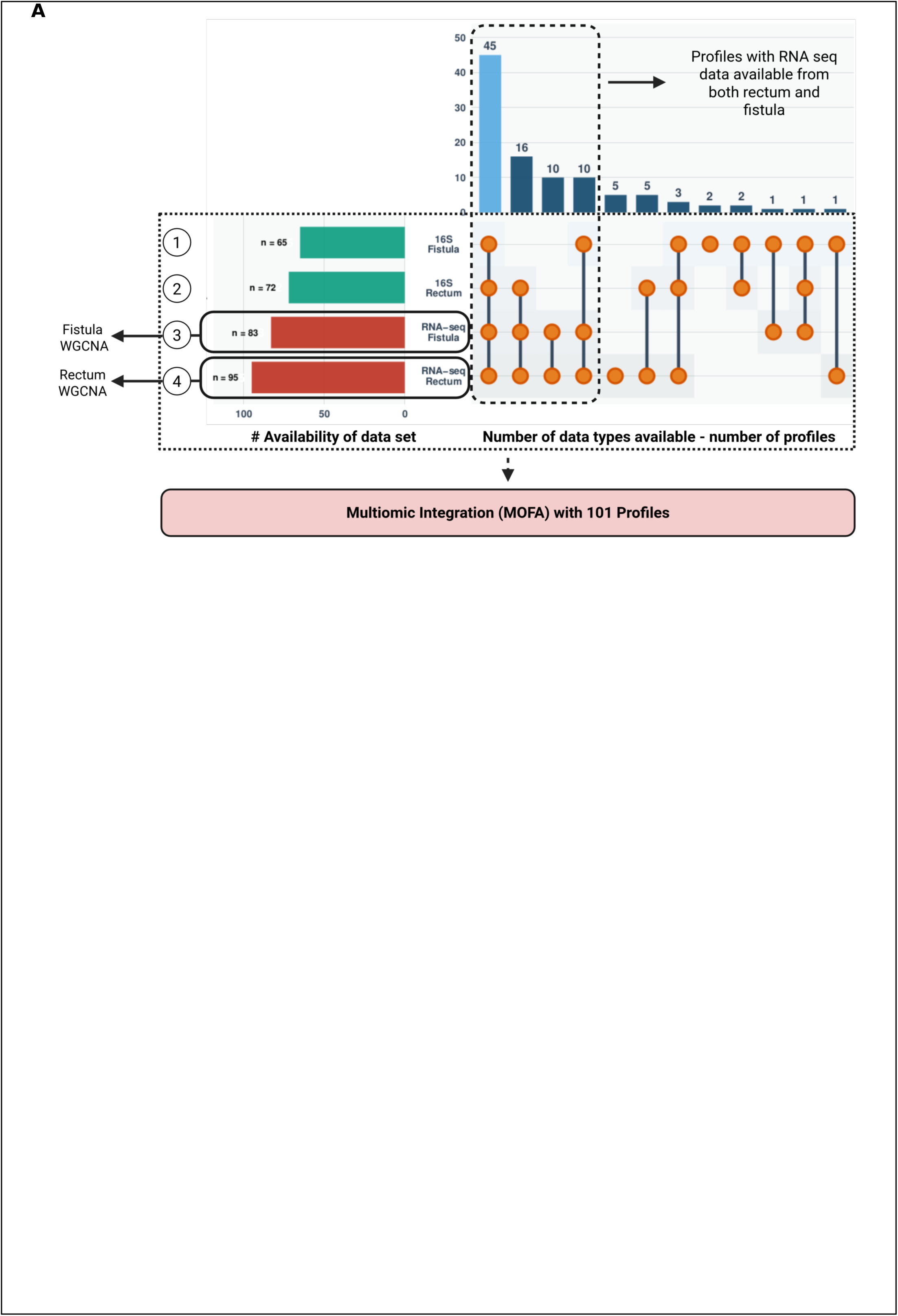
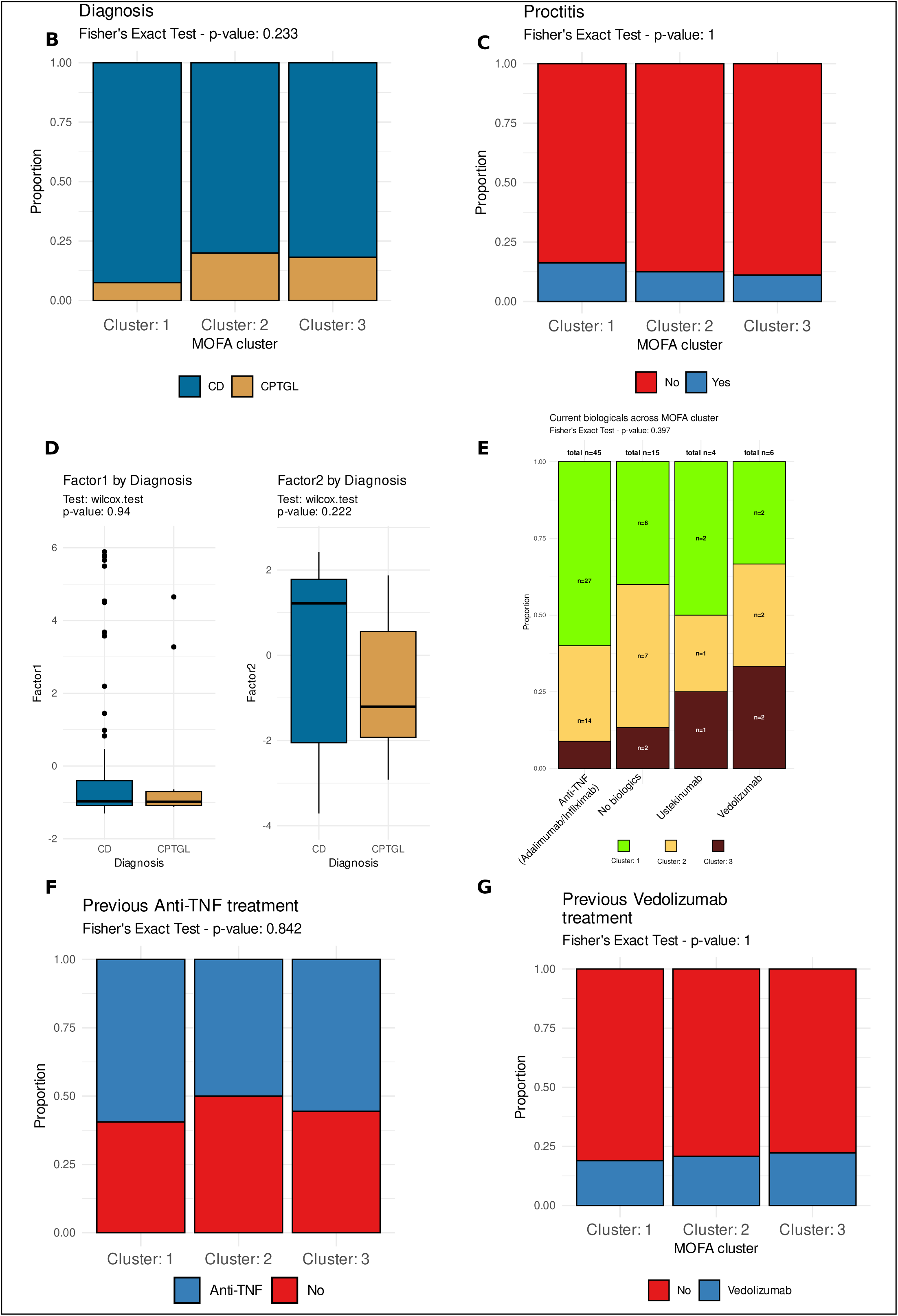
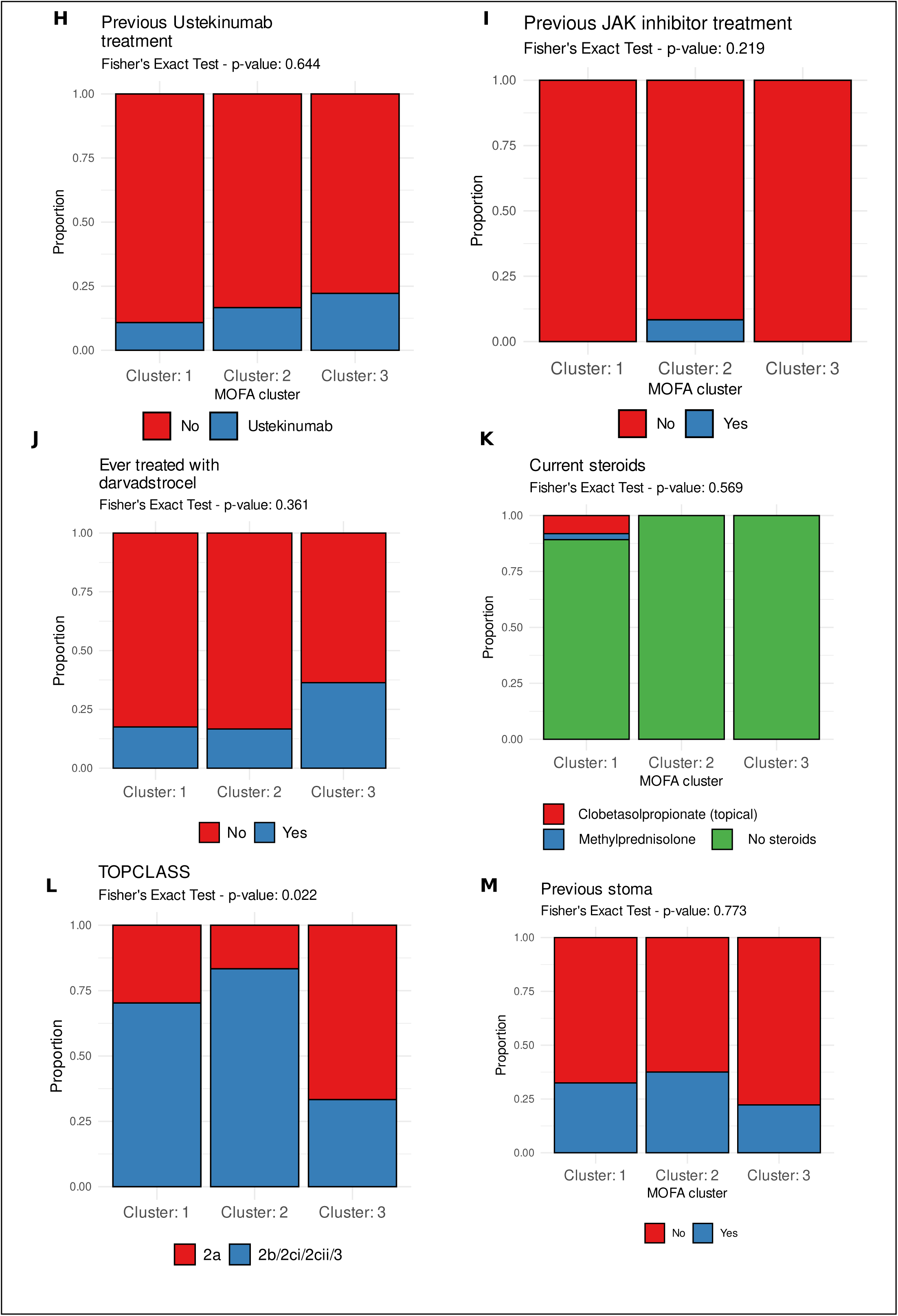
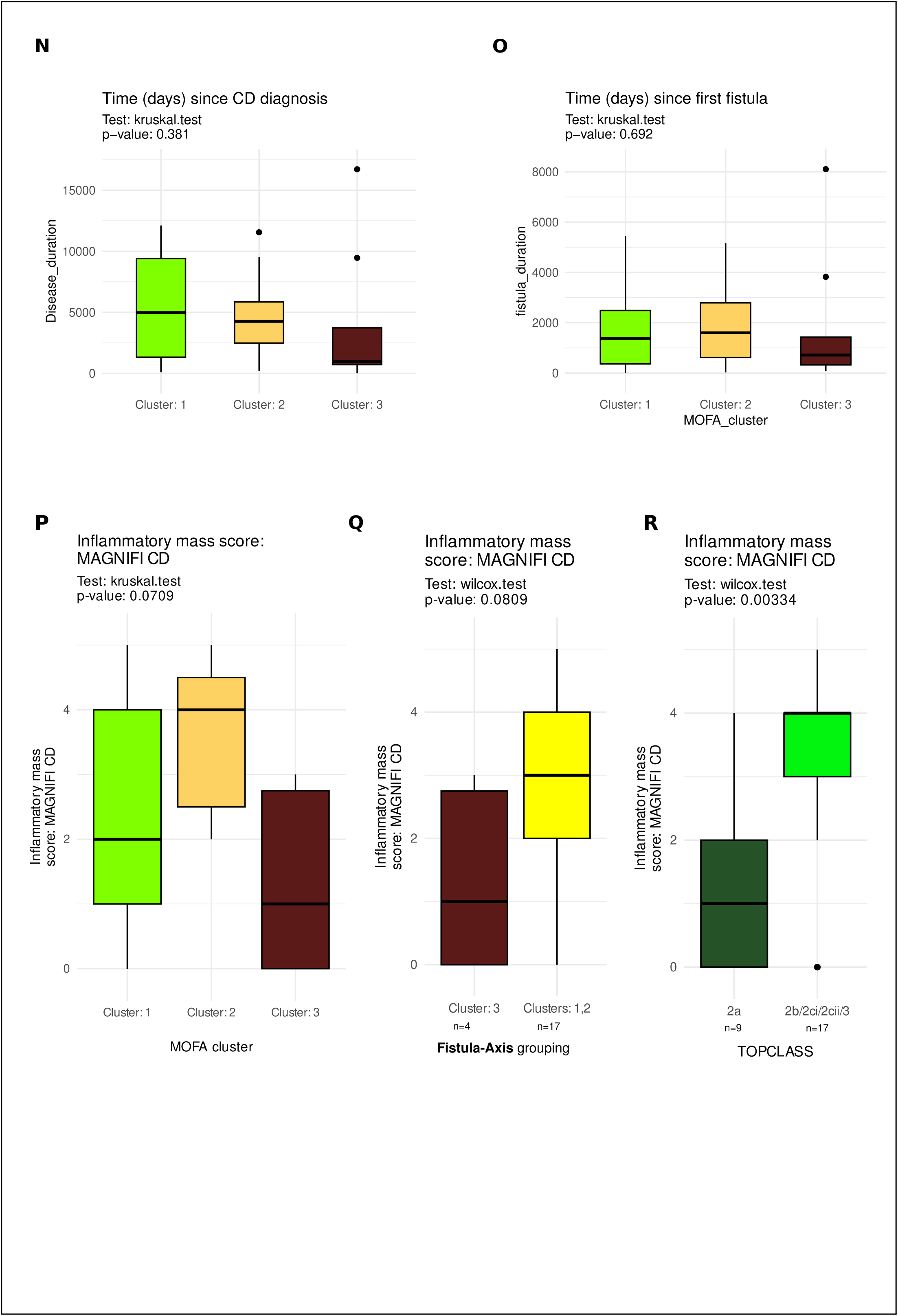
**A)** Upset plot and bar diagram showing type of ‘omic’ assay available for each profile out of 101 unique profiles that went into MOFA.**B)** Distribution of diagnosis across MOFA clusters. **C)** Distribution of proctitis status in CD patients across MOFA clusters. **D)** MOFA Factor values for Factor1 and Factor2 split by diagnosis **E-I)** Bar plots indicating previous treatments across MOFA clusters of CD patients. **J)** Bar plot depicting whether the CD patients were ever treated with darvardstrocel. **K)** Bar plot depicting status of current steroid treatment in the CD patients. **L)** TOpClass classification depicted across MOFA clusters in CD patients. **M)** Bar plot indicating previous stoma in CD patients. **N)** Disease duration (CD Diagnosis to sampling date in days) in CD patients across MOFA clusters. **O)** Time (in days) since first fistula in CD patients. **P-R)** MAGNIFI-CD inflammatory mass sub-score across MOFA clusters (**P**), fistula groups (**Q**), TOpClass categories (**R**).

**Supplementary figure 3:**
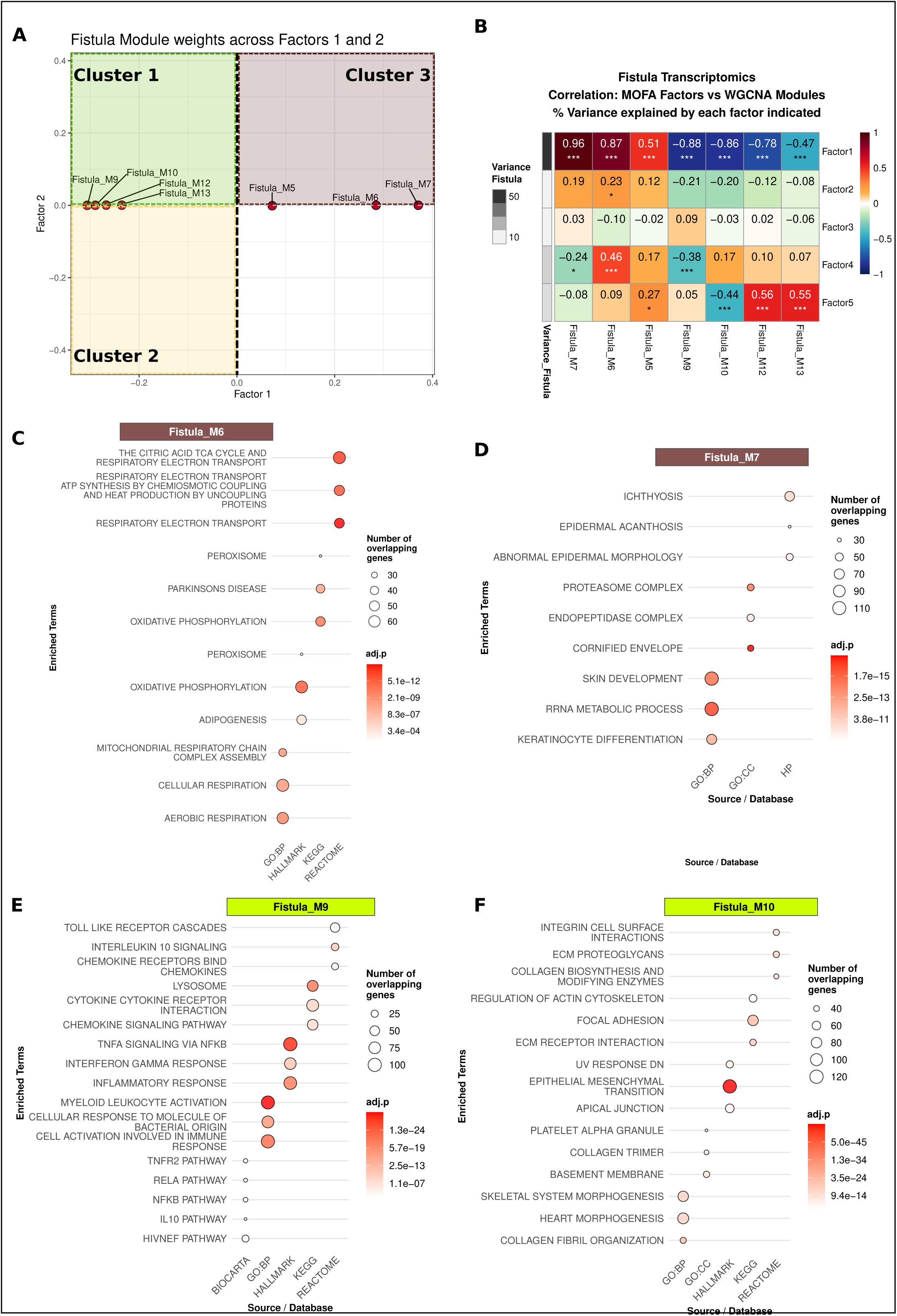
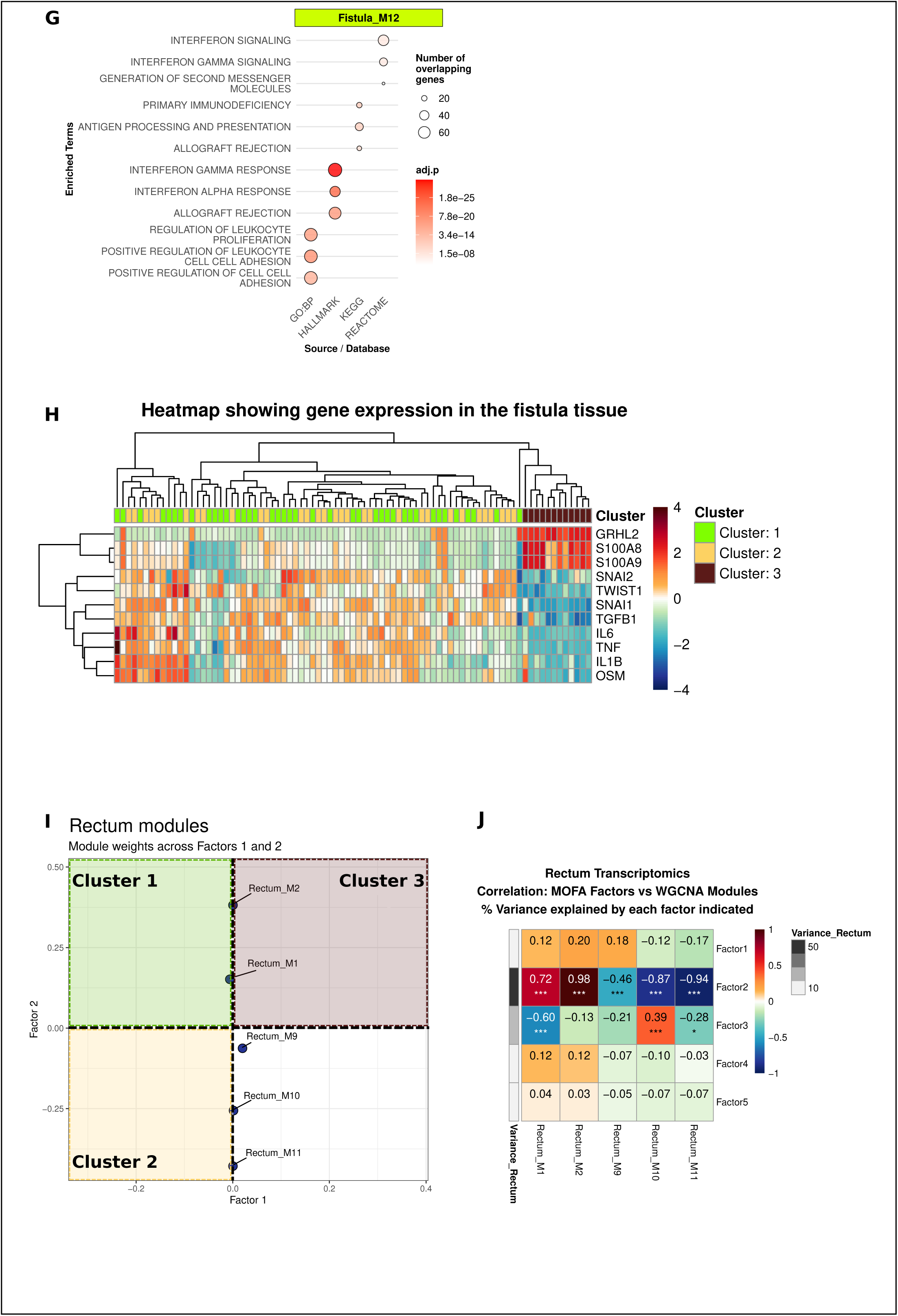
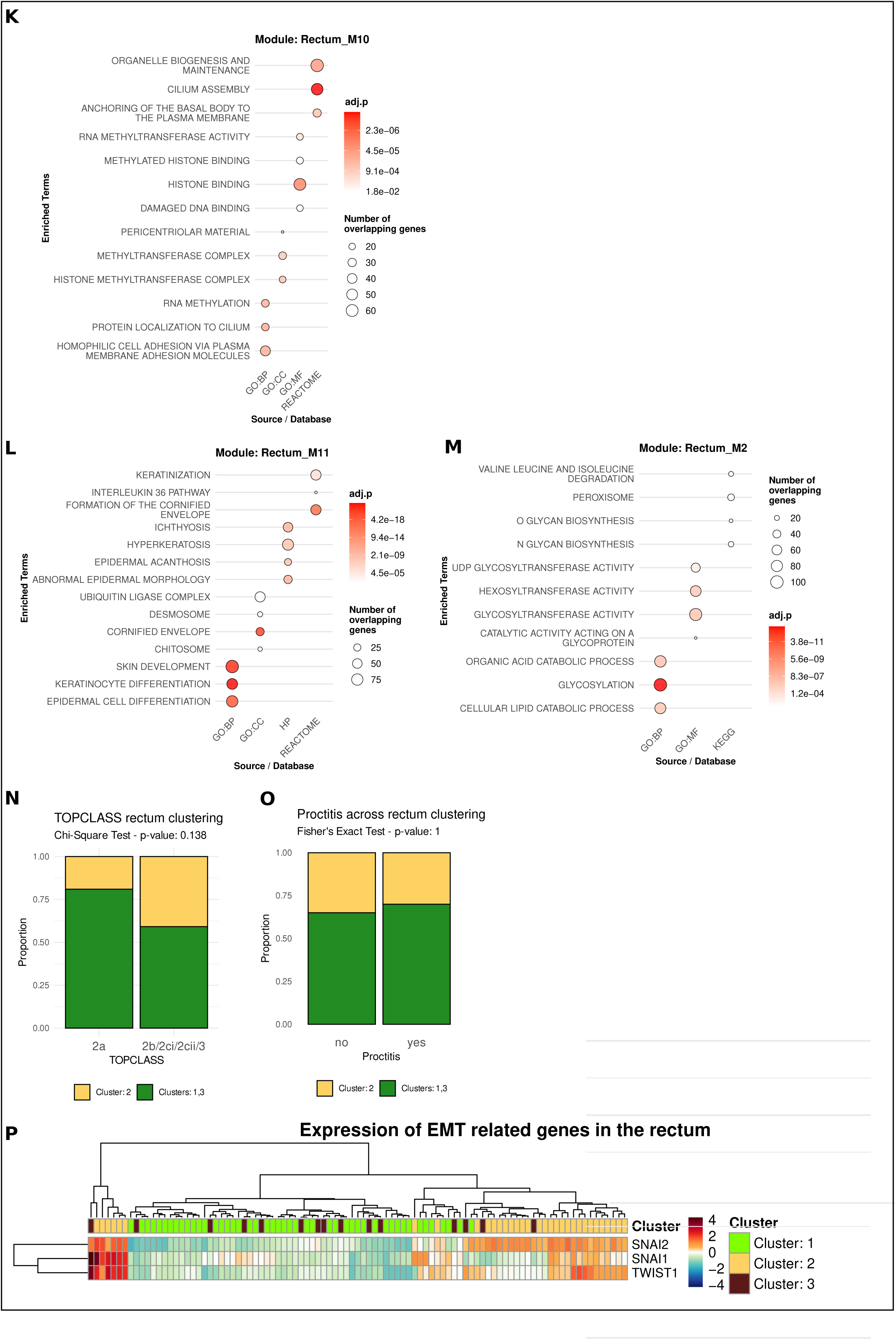
**A)** Summarized gene weights for individual WGCNA modules from fistula transcriptomics data depicted across the MOFA factor 1 and 2. Fistula modules lies horizontally along Factor 1. **B)** Correlation of fistula tissue derived WGCNA module eigengene value with MOFA factor values. Also depicted in grey the variance of fistula data explained by each factor. **C-G)** Dot plots summarizing the functional enrichment analysis of the genes in gene co-expression modules identified by WGCNA in fistula tissue. Selected terms are grouped on the x-axis by their respective database and listed on the y-axis. **H)** Normalized gene expression levels in the fistula samples across MOFA clusters (each column indicates one sample). **I)** Summarized gene weights for individual WGCNA modules from rectal transcriptomics data depicted across the MOFA factor 1 and 2. Rectal modules lies horizontally along Factor 1. **J)** Correlation of rectal tissue derived WGCNA module eigengene value with MOFA factor values. Also depicted in grey the variance of fistula data explained by each factor. **K-M**) Dot plots summarizing the functional enrichment analysis of the genes in gene co-expression modules identified by WGCNA in fistula tissue. Selected terms are grouped on the x-axis by their respective database and listed on the y-axis. (GO: Gene ontology, BP: Biological Processes, CC: Cellular components, HP: Human Phenotype (MSigDB)). **N)** TOpClass 2a vs others across the two rectum groups. **O)** Proctitis status across the two rectum groups**. P)** Heatmap showing normalized gene expression in the rectal tissue samples (each column indicates one sample).

**Supplementary figure 4:**
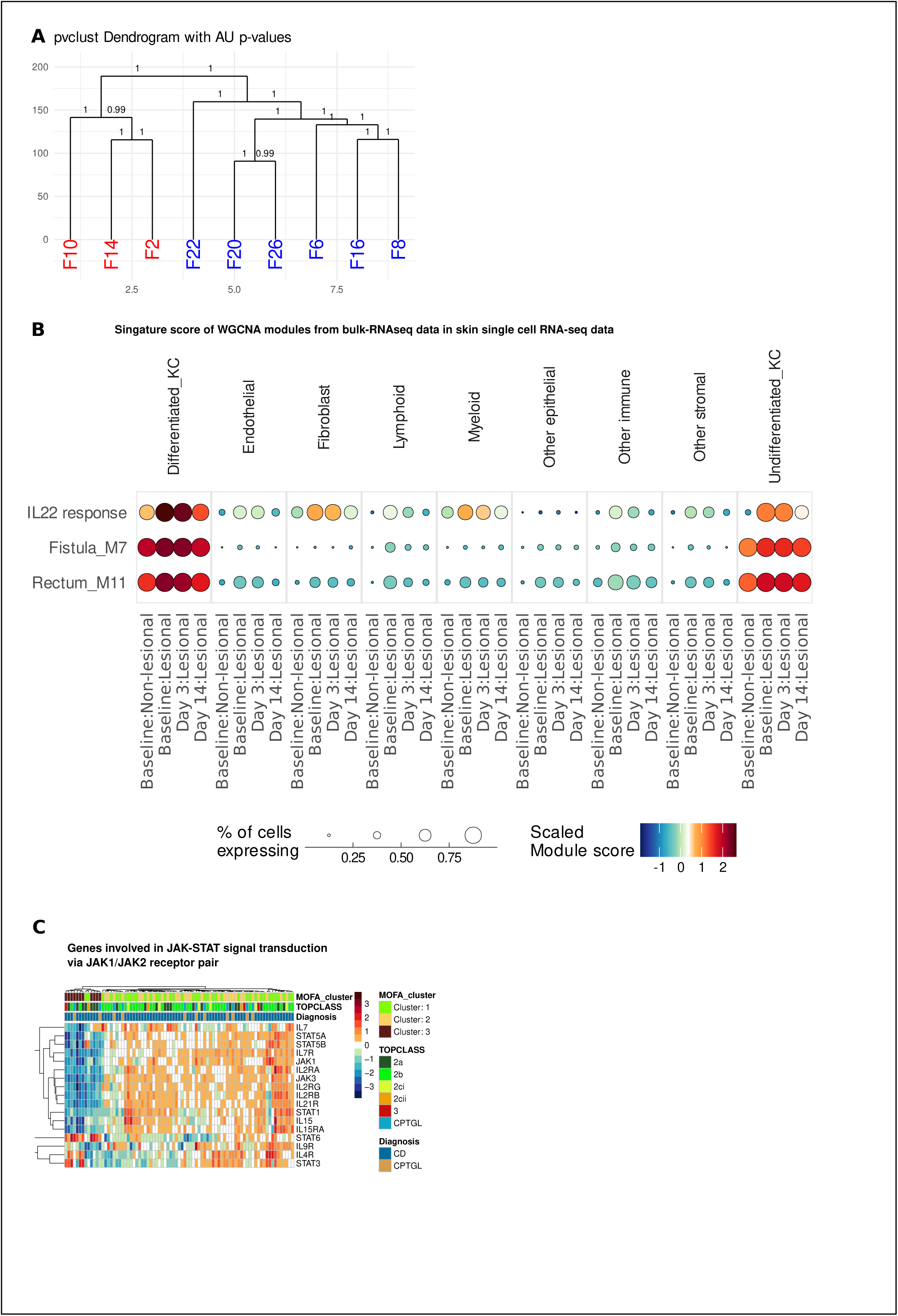
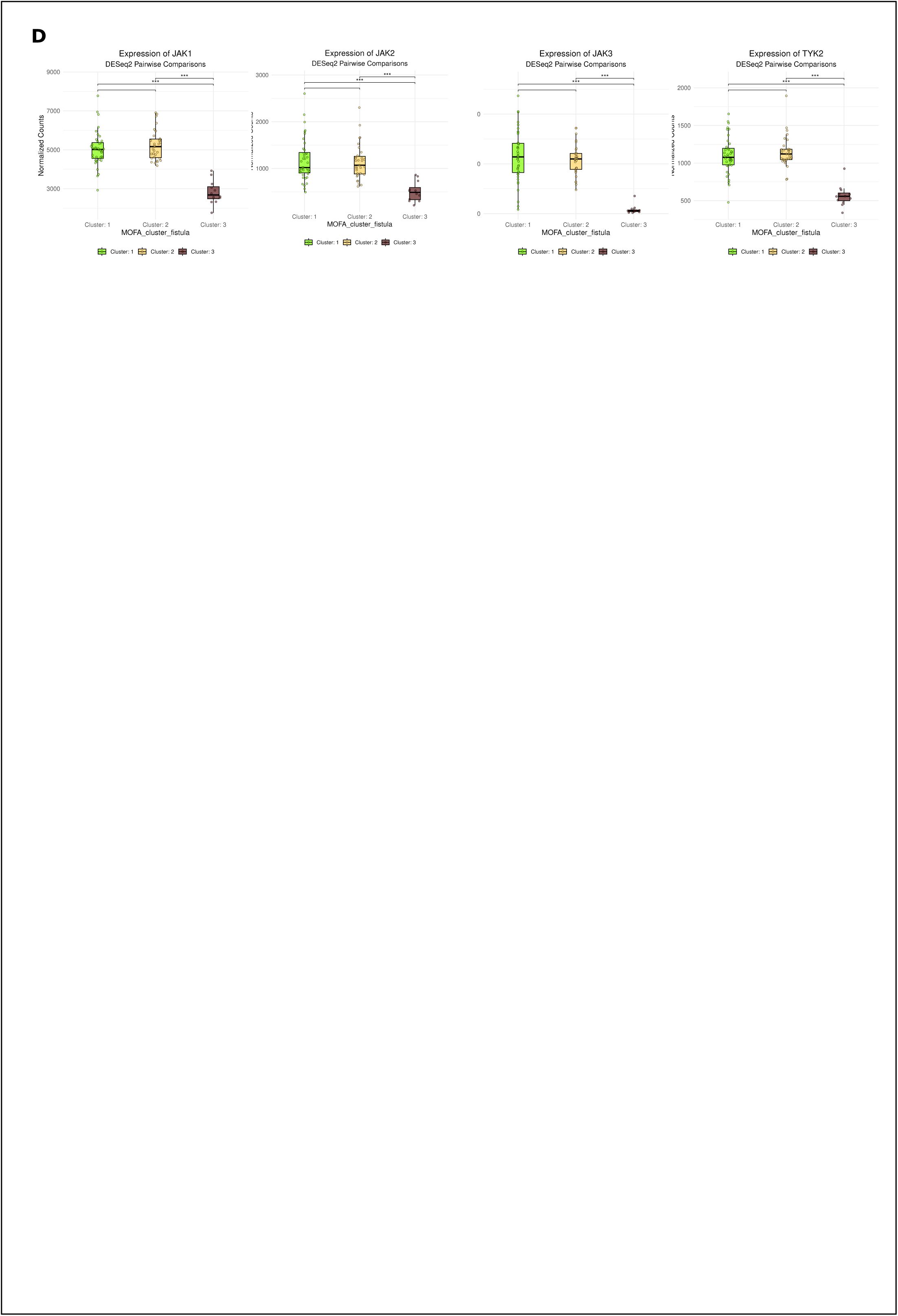
A) Hierarchical clustering dendrogram of the entire RNA-seq data from Rizzo et al., with approximately unbiased (AU) p-values annotated at each node to indicate branch confidence. B) Top hub genes (90^th^ percentile) from each of the keratinization related modules (Fistula M7, Rectum_M11) and IL22 response genes (MsigDB: Human Gene Set: ZHENG_IL22_SIGNALING_UP) was used to create a gene module score in the combined sc-RNA sequencing data from psoriasis study (Francis et. al.). Dot plot depicts the scores across different cell compartments. C) Heatmap depicting expression of genes involved in JAK-STAT signal transduction via JAK1/JAK2 receptor pair complex. D) Normalized gene expression levels in the fistula tissue across MOFA clusters.

