## Supplementary material for "UNRAVELLING A HIDDEN SUBTYPE: MULTIOMICS REVEAL PSORIASIS-LIKE SIGNATURE WITH SURGICAL RELEVANCE IN A SUBSET OF PERIANAL FISTULIZING CROHN’S DISEASE": Supp Tables

### Supplementary Tables

Table S1: Summary of Patient Characteristics (CD) at profile level

| Characteristic | Value |
|---|---|
| Patients (n) | 83 |
| Age at biopsy (y.o.; mean ± sd) | 37.2 ± 12.3 |
| Sex (M/F) | 51/32 |
| Disease duration (years.; mean ± sd) | 7.7 ± 9.3 |
| Proctitis – no. (%) | 11 (13.3%) |
| A1 below 17 y.o. – no. (%) | 14 (16.9%) |
| A2 between 17 and 40 y.o. – no. (%) | 60 (72.3%) |
| A3 above 40 y.o. – no. (%) | 9 (10.8%) |
| L1 ileal – no. (%) | 23 (27.7%) |
| L2 colonic – no. (%) | 5 (6%) |
| L3 ileocolonic – no. (%) | 36 (43.4%) |
| L3+L4 ileocolonic and upper GI – no. (%) | 13 (15.7%) |
| isolated perianal disease – no. (%) | 6 (7.2%) |
| B1 non-stricturing, non-penetrating – no. (%) | 30 (36.1%) |
| B2 stricturing – no. (%) | 14 (16.9%) |
| B3 penetrating – no. (%) | 33 (39.8%) |
| Previous anti-TNF – no. (%) | 45 (54.2%) |
| Previous vedolizumab – no. (%) | 17 (20.5%) |
| Previous ustekinumab – no. (%) | 10 (12%) |
| Ever alofisel – no. (%) | 22 (26.5%) |
| Previous stoma – no. (%) | 24 (28.9%) |
| TOPCLASS 2a | 23 (27.7%) |
| TOPCLASS 2b | 47 (56.6%) |
| TOPCLASS 2ci | 7 (8.4%) |
| TOPCLASS 2cii | 3 (3.6%) |
| TOPCLASS 3 | 3 (3.6%) |

Table S2: Summary of Patient Characteristics (CD Patients at baseline (earliest) profile)

| Characteristic | Value |
|---|---|
| Patients (n) | 48 |
| Age at biopsy (y.o.; mean ± sd) | 36.3 ± 13.8 |
| Sex (M/F) | 25/23 |
| Disease duration (years.; mean ± sd) | 5.8 ± 8.5 |
| Ethnicity:Caucasian | 100% |
| Proctitis – no. (%) | 10 (20.8%) |
| A1 below 17 y.o. – no. (%) | 6 (12.5%) |
| A2 between 17 and 40 y.o. – no. (%) | 36 (75%) |
| A3 above 40 y.o. – no. (%) | 6 (12.5%) |
| L1 ileal – no. (%) | 7 (14.6%) |
| L2 colonic – no. (%) | 4 (8.3%) |
| L3 ileocolonic – no. (%) | 29 (60.4%) |
| L3+L4 ileocolonic and upper GI – no. (%) | 5 (10.4%) |
| isolated perianal disease – no. (%) | 3 (6.2%) |
| B1 non-stricturing, non-penetrating – no. (%) | 19 (39.6%) |
| B2 stricturing – no. (%) | 7 (14.6%) |
| B3 penetrating – no. (%) | 19 (39.6%) |
| Previous anti-TNF – no. (%) | 26 (54.2%) |
| Previous vedolizumab – no. (%) | 12 (25%) |
| Previous ustekinumab – no. (%) | 8 (16.7%) |
| Ever alofisel – no. (%) | 14 (29.2%) |
| Previous stoma – no. (%) | 13 (27.1%) |
| TOPCLASS 2a | 19 (39.6%) |
| TOPCLASS 2b | 19 (39.6%) |
| TOPCLASS 2ci | 5 (10.4%) |
| TOPCLASS 2cii | 3 (6.2%) |
| TOPCLASS 3 | 2 (4.2%) |

Table S3: Summary of Profile Characteristics (CPTGL Patients at Profile level)

| Characteristic | Value |
|---|---|
| Patients (n) | 18 |
| Age at biopsy (y.o.; mean ± sd) | 56 ± 15.5 |
| Sex (M/F) | 14/4 |
| No Antibiotics (no. (%)) | 16 (88.9%) |

Table S4: Summary of Patient Characteristics (CPTGL - at baseline (earliest) level)

| Characteristic | Value |
|---|---|
| Patients (n) | 15 |
| Age at biopsy (y.o.; mean ± sd) | 56 ± 14.4 |
| Sex (M/F) | 12/3 |
| Ethnicity:Caucasian | 100% |
| No Antibiotics (no. (%)) | 13 (86.7%) |

Table S5: Summary of MRI availability across MOFA clusters and TOpClass categories

| | Cluster 1 | Cluster 2 | Cluster 3 | **Total** |
|---|---|---|---|---|
| TOpClass 2a | 4 | 1 | 4 | **9** |
| TOpClass 2b | 6 | 5 | 0 | **11** |
| TOpClass 2cii | 0 | 1 | 0 | **1** |
| **Total** | **10** | **7** | **4** | **21** |
